# Continuous Integration and Software Quality in Scientific Software: A Large-Scale Empirical Analysis of GitHub Repositories

**DOI:** 10.64898/2026.10.01.755028

**Authors:** Walker J. Compton Mellon, Makena Sy, Navya Gangrade, Tobias Jakobi

## Abstract

Bioinformatics software plays a central role in biological research, yet many academic projects lack robust testing and continuous integration (CI) practices. This absence can compromise reproducibility, accessibility, and long-term maintainability. In this study, we conduct an large-scale empirical analysis of GitHub repositories spanning bioinformatics and other scientific software domains to examine the relationship between CI, software testing, and software quality. Repository discovery was performed using domain-specific search terms via GitHub’s Application Programming Interface (API). This was followed by a layered data extraction pipeline, which included a Large Language Model (LLM) validation step, that connected repositories to published works, institutions, authors, funding, and citation counts. By integrating repository activity, community engagement, and backlog health into a composite Repository Quality Index (RQI), we find that the presence of CI and software tests is significantly associated with higher repository quality. Additionally, repositories with higher academic citation counts tend to cluster in higher RQI tiers, suggesting that software quality, which is driven by proper testing and integration, is associated with academic impact and funding status. These results demonstrate that adopting industry-standard development practices, such as CI and software testing, is a tractable path toward more sustainable and impactful academic software and funding opportunities.

## Introduction

Bioinformatics software is crucial to modern biological research, facilitating significant advances in data analysis and discovery ^[1]^. Open-source bioinformatics software, in particular, has been instrumental in driving scientific discovery from biological data ^[2]^. Because these tools directly inform scientific conclusions, their outputs must be reproducible across different computational environments ^[3]^. Nevertheless, despite their critical contributions, such tools often exhibit inconsistencies in accessibility, reproducibility, and long-term support ^[4]^.

The ecosystem that bioinformatics tools build themselves upon is fragile and ever-changing. This volatility is driven by changes across operating systems, hardware architectures, and most importantly, the dependency stack of software libraries and packages. Although many of these changes and the failures caused by them are hard to foresee, developers and maintainers can utilize continuous integration (CI) and testing protocols to ensure that bioinformatics software remains functional, reproducible, and accessible as computational environments and infrastructure evolve. Despite the widespread recognition of these challenges, there is limited empirical analysis of unified software quality indicators that could drive improvements in scientific software development practices ^[5,6]^. Previous studies utilize a similar API based repository data collection approach, but focus on direct software performance and internal development benefit. This study addresses that gap by conducting a large-scale empirical analysis of GitHub repositories to examine the relationship between CI, software tests, software quality, and academic performance.

Open-source software is deeply embedded in today’s computational pipelines, web infrastructure, and research tools. Development of these projects is typically distributed across a wide range of contributors, including individual volunteers, research groups, and organizations, whose contributions span code, documentation, bug reports, and feature requests. Managing and integrating this breadth of contributions places an ongoing maintenance burden on project developers, who may range from a single volunteer maintainer to a team of dedicated engineers. This wide variation in resources, development practices, and contributor base produces substantial variance in maintenance approaches, which can deeply impact the long-term sustainability of a project.

Like all software systems, bioinformatics tools are subject to a lifecycle shaped by the surrounding computational ecosystem. Deprecation is rarely abrupt; unsupported tools can gradually become unusable as operating systems, programming languages, and external dependencies evolve around them. This challenge is particularly pronounced in bioinformatics, where software development is often tied directly to academic research projects. Many tools are built by early-career researchers, sometimes with limited training in classical software development concepts, whose priority is scientific discovery rather than long-term software sustainability. As these developers transition to new projects or institutions, active maintenance frequently ceases, leaving tools to grow incompatible with modern systems and placing substantial bodies of research output at risk. Proper end-of-life (EOL) planning can turn software deprecation from an abrupt collapse into a managed transition. For example, when phasing out Python 2 in favor of Python 3, the Python developers gave the community advance notice and an initial defined transition window of 7 years before fully retiring Python 2 ^[7]^. Frameworks that support software testing, reproducibility, maintainability, and longevity can help mitigate these risks by reducing the technical burden of software handoff and extending the functional lifespan of critical tools.

To address the challenges posed by diverse operating systems, programming languages, and dependencies, modern software engineering relies on integration testing and CI frameworks ^[8]^. Integration testing evaluates how well individual software components interact with one another and with external systems, such as different operating environments and libraries. CI embeds these tests directly into the development cycle, ensuring that every new code change is automatically validated for compatibility and functionality. This process helps maintain stable, reproducible, and reliable software throughout its evolution.

Despite the availability of these practices, users of bioinformatics software frequently encounter challenges stemming from the complex network of interrelated packages, libraries, and external dependencies that most workflows rely on. Although these dependencies often function reliably within controlled development environments, their behavior in real-world applications can be inconsistent. As the broader software ecosystem evolves, deprecation and breaking changes can quietly accumulate until a once-functional tool becomes unreliable or entirely unusable. Continuously validating that these components work together ensures that such issues are detected and resolved promptly, rather than discovered by end users and left unresolved.

In academic research, CI and tests often take a backseat to the central development focus, despite their clear benefits. In industry, code and testing go hand in hand, so much so that shipping untested code would be considered a liability, since test coverage is a standard expectation of the development process ^[9]^. Additionally, industry developers are accountable to stakeholders who require varying levels of software quality, driven by the financial incentives available within the industry. Academic developers face different requirements. Rather than stable delivery or financial gain, these developers focus on creating novel software that drives scientific discovery. Software compatibility and portability may not fall into focus, since publications and citations are more relevant measures of success in academia. This study shows that investing time and effort into sustainable software development practices can potentially boost academic outcomes for projects.

## Methods

### Repository Discovery and Filtering

To identify trends across GitHub repositories, we first collected metadata from a large and diverse set of repositories. An initial step in the process was determining which repositories to include in the analysis. We began by defining a set of broad, representative search terms corresponding to relevant scientific and software development domains. These terms included bioinformatics, astrophysics, open source, software engineering, and image recognition.

These search terms were selected to capture a diverse range of scientific and non-scientific software projects, enabling comparative analysis of repository maintenance practices and software development trends. Using these terms, we queried GitHub’s REST API to assemble a large, pre-filtered collection of 381,078 repositories (Supplementary Table ST1).

The collection of repositories was then used to query GitHub’s API to collect detailed data. First, general information about the repositories, such as star, commit, and issue counts, were queried from GitHub’s GraphQL API. This GraphQL query was also used to detect GitHub workflow files and other CI configuration files, which are used for CI. If these files were present the repository would be marked as having CI. Next, the GitHub REST API was called to retrieve a recursive file tree. Keywords indicating software tests (e.g., “test”, “conftest.py”, “makefile.py”) were searched for within the file tree. Repositories with matching paths were then marked as having software tests. The REST API also was also used to retrieve commit and issue counts, history, and information.

Citation counts were then gathered for each repository to track the academic success of these software tools. This was achieved through a layered extraction pipeline that uses information within the repository to find a valid reference in Web of Science (WoS). WoS is a database that indexes citation information, publications, and author and institutional information. The pipeline operated as a priority chain across four layers, terminating at the first successful match. The first layer checks for the presence of a structured citation file, such as CITATION.cff, CITATION.bib, which are suggested and supported by GitHub to enable clear project citations, or R-package conventions like inst/CITATION. Digital Object Identifiers (DOIs), centralized identifiers managed by the DOI Foundation, along with paper titles and author names, were parsed from these files according to their format. DOIs associated with academic data repositories (e.g., Zenodo, Figshare, Open Science Framework) were excluded as non-paper identifiers, as these identify datasets or software archives rather than academic papers. If a valid DOI or title was extracted, it was queried against WoS. If the publication was identified but not indexed in WoS, the repository was recorded as such, and no further layers were attempted. If no citation file was present or the citation file did not yield a WoS result, the pipeline scanned the README for PubMed URLs, resolving them to DOIs via the NCBI E-utilities API before querying WoS ^[10]^. If no PubMed identifier was found, the pipeline scanned the README for DOIs matching known academic publisher prefixes and queried them directly. If all prior layers failed, a local large language model (LLM, Gemma 4 26b at temperature 0.0) was prompted with the repository name, tool name, pre-scan signals detected via regex (including BibTeX blocks, citation language, journal mentions, and arXiv IDs), and the first 4,000 characters of the README content. The model returned structured JSON containing extracted DOIs, PubMed IDs, author names, and a composite search string. This was then used to construct a list of layered WoS queries that included DOI lookup, author name combined with tool name, search title, tool name combined with subtitle, and explicit title. Any match returned was subject to a validation step, in which the LLM was called to confirm that the retrieved paper described the tool itself, rather than a paper that merely cites or uses the tool or the foundational findings underpinning it.

### Repository Quality

Repository quality cannot be directly measured and was therefore inferred from three observable dimensions: backlog health, recent activity, and popularity. Backlog health reflects how effectively a repository addresses reported issues. A lower median of issue resolution time reflects sustained developer attention, consistent bug fixes, and quick feature request fulfillment. Recent activity was measured as the time elapsed between data collection and the most recent commit, serving as an indicator of ongoing development and maintenance, including compatibility with evolving software dependencies and computational environments. Popularity was defined as a composite measure derived from a repository’s stars, watchers, and forks, capturing the extent of community adoption and engagement.

### Quality Assessment and Ranking

Before repositories were compared and analyzed, they underwent additional filtering. Repositories with fewer than ten commits and ten total issues were removed to ensure that the projects being analyzed were legitimate development projects. From here, each quality dimension (backlog health, popularity, and recency) was scored by anchoring specific indicator values to fixed ratings, then applying linear interpolation to produce a continuous score. For recency, a commit on the day of data collection scored a 5, easing down through two weeks, one month, two months, three months, six months, to 1.3 at a year and 0.0 at two years and longer.

For backlog health, the median of the length of time issues stayed open was calculated then subjected to the anchoring system. A median of issue closures on the same day was scored at 5.0, easing down through one week, two weeks, one month, two months, four months, and to 1.0 at a year. Lastly, popularity was calculated from the log_10_ transformation of the sum of stars, forks, and watchers. Each order of magnitude of popularity is correlated with rating, ranging from a rating of a 1.0 for a total of zero popularity metrics to 5.0 for a total of 9,999 or more. The three anchor-based dimensional scores were then averaged to produce the Repository Quality Index (RQI, Supplementary Table ST2).

### Tracking Funding for Project Organizations

To track relationships between software development practices and project funding, we used a multi-stage pipeline that linked GitHub repositories to grant funding records across multiple agencies via the OpenAlex database and the NIH RePORTER API. OpenAlex is a digital catalog of scholarly data, including information on academic works, institutions, and research funding ^[11]^. Each repository was associated with a publication DOI, which was resolved to a PubMed ID (PMID) via the NCBI ID Converter, eSearch, and Europe PMC APIs. Grant information was supplied via a local snapshot of the OpenAlex database (snapshot dated June 26, 2026), enabling bulk lookup of funding status and amounts across multiple agencies, including NIH, NSF, DOE, UKRI, ERC, and others. NIH grants were additionally enriched through the NIH RePORTER API to retrieve direct costs and activity codes not present in the OpenAlex schema. For each match between a repository and a grant, a locally served LLM (Gemma 4 26b) classified whether the grant directly funded the development of the tool, using the grant abstract, README, author affiliations, and PI names as context. Center grants and training grants were subject to additional constraints to prevent institutional affiliation alone from being treated as evidence of direct funding. This ensures that only projects directly funded by an agency are retained in the dataset, while those that simply cited or utilized the tool are excluded.

## Results

A total of 13,670 repositories met the activity thresholds across the software domains, which was a minimal commit count of 10 and an issue count of 10. Across the datasets, 62.8% of repositories had CI configured, and 87.3% had testing, with 7.5% lacking both (Supplementary Tables ST3-ST4).

### Quantifying and Comparing Software Quality across Domains

To quantify software quality across domains, RQI was computed for all 13,670 repositories. This yielded an overall mean of 2.88 (median 2.71) on the 0-5 scale. The components of RQI were standardized to fit a 0-100% scale within a radar plot (Fig. 1). Before hypothesis testing, the normality of the RQI distribution was assessed (Supplementary Figs. S1-S3, Supplementary Tables ST5-ST6). A one-way ANOVA conducted across all groups suggested significant differences in RQI across domains (p<0.001, F(4, 13665) = 179; Supplementary Tables ST7-ST8). Projects within the astrophysics domain had the highest RQI (2.97), followed by open-source (2.96), bioinformatics (2.79), image recognition (2.56), and software engineering (2.22) (Table 1). Tukey post-hoc comparisons confirmed that bioinformatics scored significantly lower than astrophysics (p=0.02) and open-source (p<0.0001), and significantly higher than image recognition (p=0.03) and software engineering (p<0.0001, Fig. S4; Supplementary Table ST9). These findings suggest that bioinformatics occupies a middle tier of repository quality across scientific domains. Examining the individual RQI components can help explain this positioning. Bioinformatics had the second-highest recency score (88.1), indicating active and recent development, but had the lowest mean backlog health (0.22) across all domains, suggesting that while bioinformatics repositories are actively committed to, they are the slowest at resolving issues. Bioinformatics ranked third in mean popularity (3.74), which reflects the niche subject matter of many bioinformatics packages and software.

**Fig 1.**
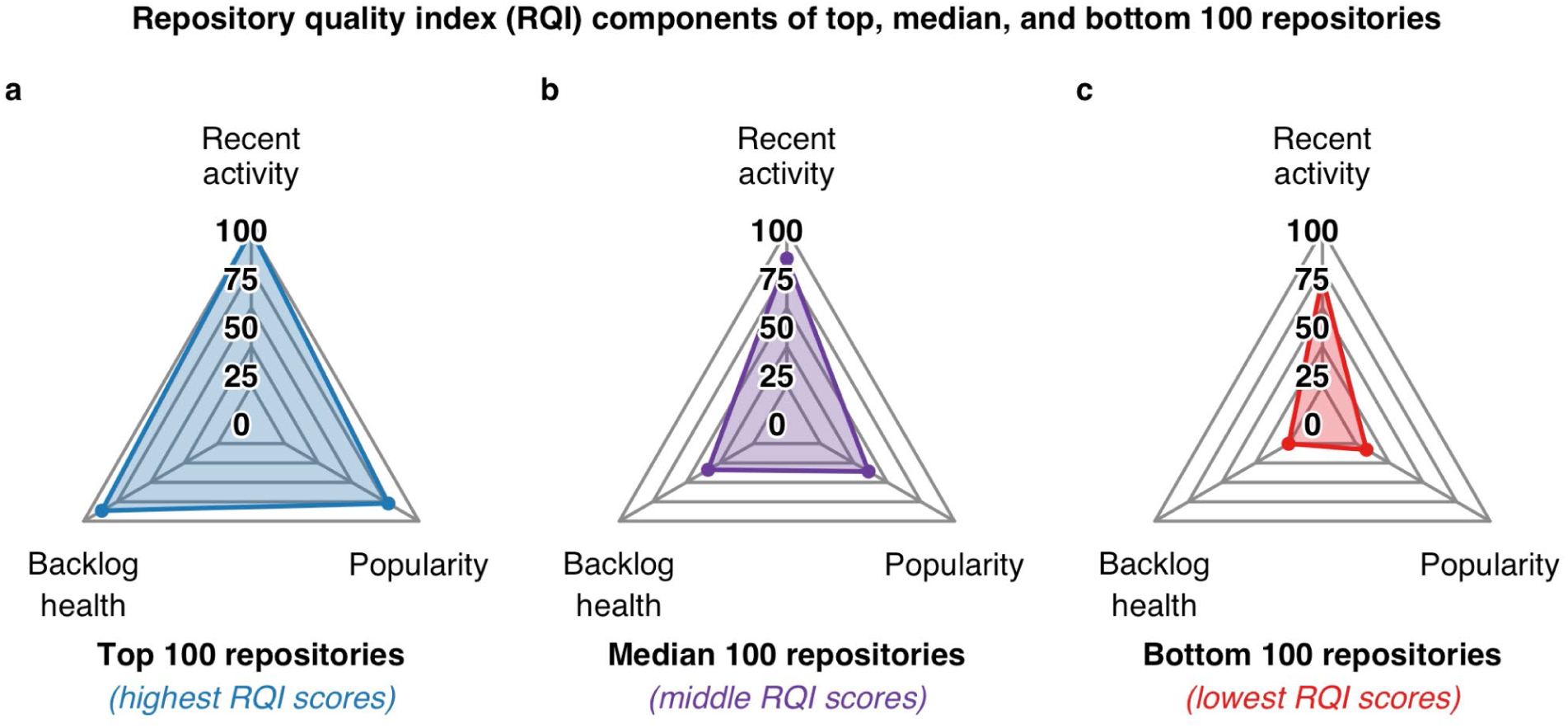
Radar Plots for RQI Breakdown of Top Repositories. Average metric profile for the top (**a**), median (**b**), and bottom (**c**) 100 repositories across all five domains. Each plot shows each aspect of the RQI (backlog health, recent activity, and popularity) that is normalized to fit the plot. Values are normalized between 0 and 100 from the mean values.

**Table 1.**
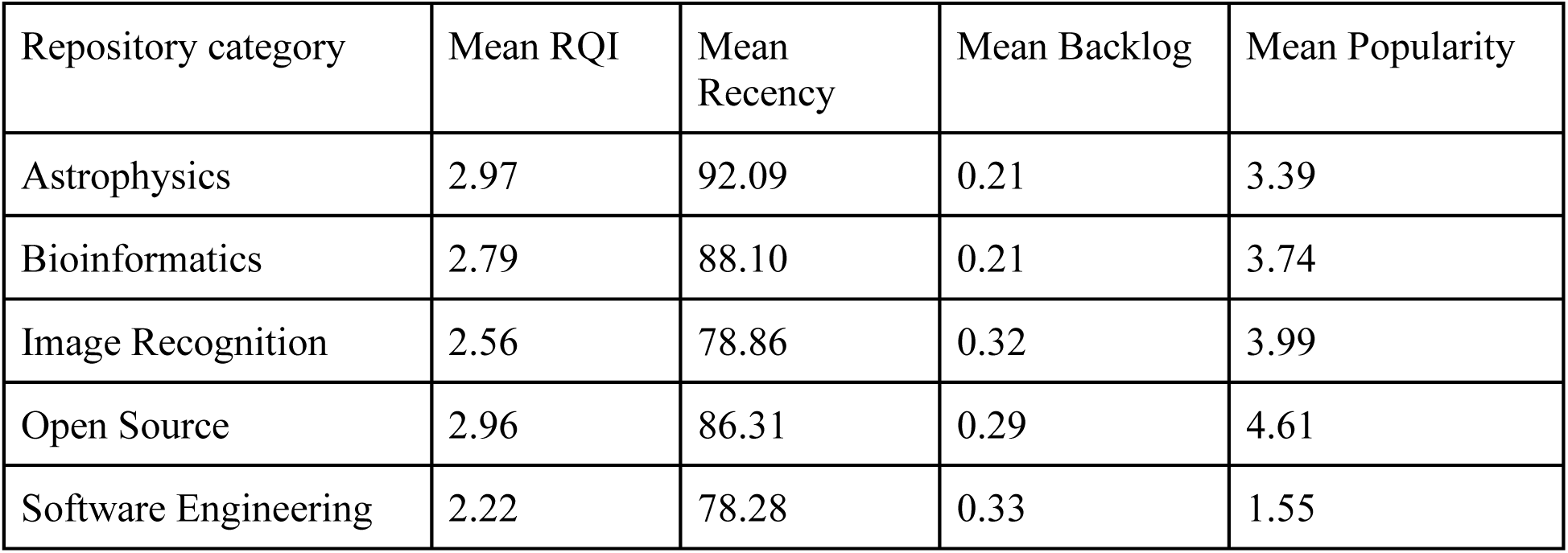
Mean Scores by Repository Category. (mean RQI, recency, backlog, popularity)

The repositories that fall within the top 100 and median 100 repositories have overall high RQI Scores (averages of 100 and 81.8) for recent activity, and the bottom 100 with an average of 69.0 (Supplementary Table ST10). On average, in relation to backlog health, the top 100 repositories resolve issues roughly 2.5 times faster than the median repositories (33.8) and nearly 432 times faster than the bottom 100 (0.2). Popularity across the top 100 repositories sat at an average of 77.2, the median at 36.0, and the bottom at 7.9. The top, median, and bottom values for the bioinformatics domain followed a similar pattern. Recent activity from top to bottom was 99.8, 89.0, and 74.9, respectively. Backlog health for top, median, and bottom was 39.8, 25.7, and 0.8. Popularity did not range as much with the top 100 having a score of 43.4, the median with 30.7 and the bottom with 21.1 (Supplementary Table ST11, Supplementary Fig. S5). The average profile per domain revealed that astrophysics had the highest score for recent activity (92.1), software engineering had the highest score for backlog health (33.3), and open-source software had the highest score for popularity (35.2) (Supplementary Fig. S6, Supplementary Table ST12). Comparing the top and bottom repositories within each domain shows how the characteristics of RQI components separate high-quality repositories from low-quality repositories (Supplementary Fig. S7, Supplementary Table ST13). For example, the bottom 20 astrophysics repositories have larger popularity (20.3) and recent activity scores (81.5) than all of the other bottom 20 repositories in every other domain.

### Trends in Software Quality

CI and testing prevalence increased consistently with repository quality (Fig. 2, Supplementary Table ST14). Linear regression on RQI-binned prevalence estimates revealed a strong positive association between RQI and CI adoption (slope = 0.15 per RQI unit, R² = 0.94, p < 0.001, Fig. 2), while testing prevalence showed a weaker but still significant relationship (slope = 0.04, R² = 0.69, p < 0.001, Fig. 2). The steeper slope and greater explained variance for CI suggest that it is a stronger predictor of repository quality than testing alone. This pattern persisted when repositories were binned into low (RQI ≤ 2), medium (2 < RQI < 4), and high (RQI ≥ 4) quality groups, with quality groups significantly differing from each other (Supplementary Fig. S8, Supplementary Table ST15). CI prevalence rose from 45.9% among low-quality repositories to 89.3% among high-quality repositories, while testing prevalence increased only modestly from 84.1% to 94.9% across the same range. The narrow spread in testing prevalence across quality tiers indicates that while general testing is widespread even among lower-quality projects, it is not sufficient on its own to distinguish between high- and low-quality repositories. Welch two-sample t-tests confirmed that repositories with CI had significantly higher mean RQI than those without (3.09 vs. 2.51, p < 0.001), as did repositories with software tests (3.91 vs. 2.62, p < 0.001; Supplementary Fig. S9, Supplementary Table ST16). However, the effect size for CI was substantially larger (∼0.57) than that for testing (∼0.29), reinforcing that CI is a more meaningful indicator of repository quality. This pattern held within each software domain. Repositories with CI and software tests had significantly higher mean RQIs, except for testing in the software engineering domain (Supplementary Table ST17, Supplementary Figs. S10-S11). These findings suggest that while software testing is common practice even among lower-quality projects, CI represents a higher standard of software development that more consistently separates poorly maintained projects from high-performing tools with sustained software support.

**Fig 2.**
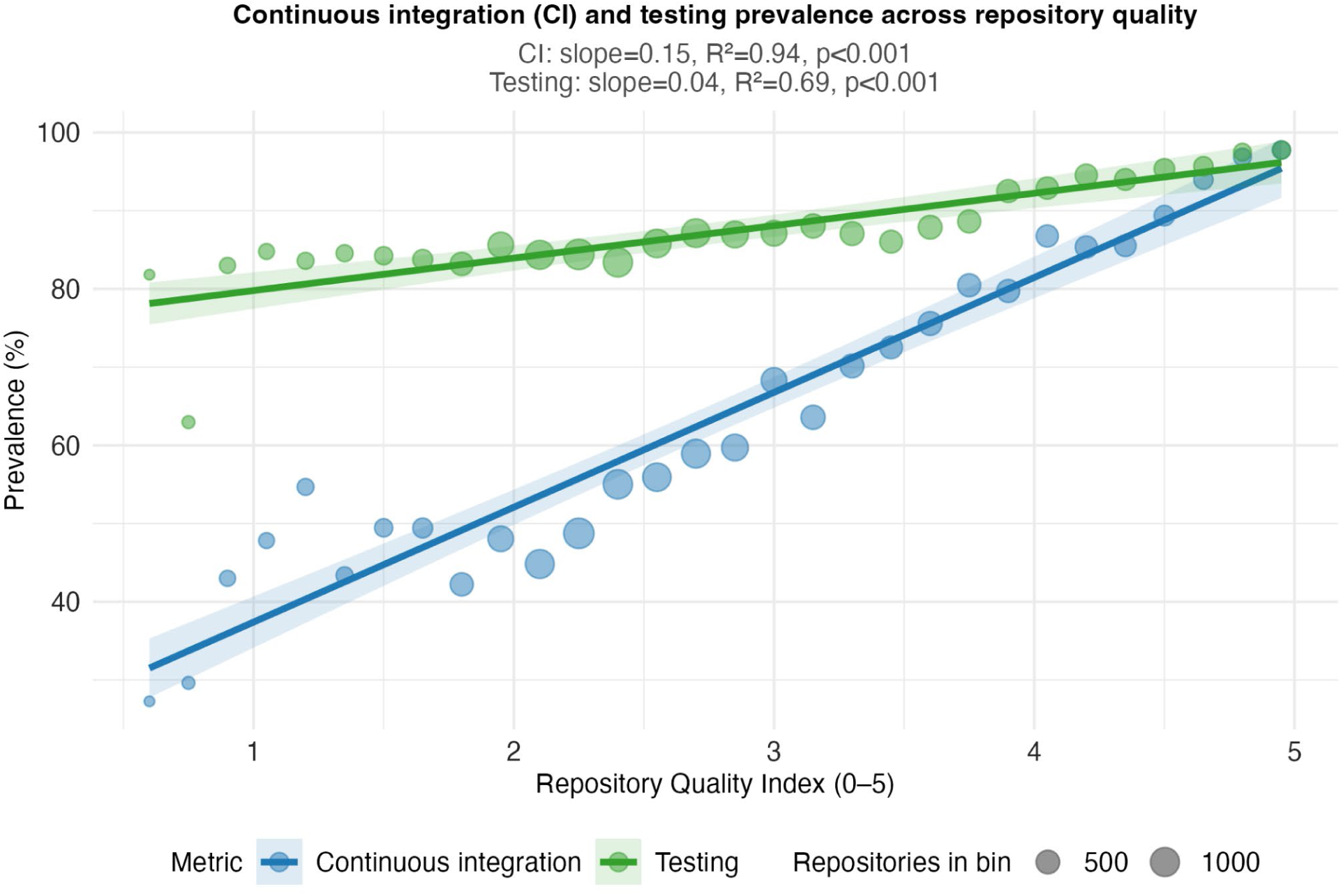
Continuous Integration/Testing Prevalence in RQI Quality Groups. Repositories are binned into groups at each 0.5 increment of RQI. Prevalence is calculated as the proportion of the number of repositories that adopt CI/Testing divided by the total number of repositories within the quality group.

### Overview of Software Languages within GitHub Repositories

Overall, the top 3 programming languages with at least one representative file within a repository were Javascript (n = 6059), Shell (n = 5501), and Python (n = 5152; Supplementary Table ST18). The top 3 most frequent primary languages were Python (n = 2572), Javascript (n = 1635), and Typescript (n =1505; Supplementary Table ST19). The 3 primary software languages with the highest mean RQI (MDX, Rust, and GO) had CI coverages of 85.7%, 88.5%, and 86.9% and testing coverages of 96.4%, 90.7%, and 91.8%, respectively (Supplementary Table ST20). Within each language, repositories with CI had a higher mean RQI than those without CI. The same was held for testing for 24 out of 28 of the languages with an available comparison, with the exceptions being MDX, Javascript, Dockerfile, and Fortran (Supplementary Table ST21, Supplementary Fig. S12-S14).

Within bioinformatics, the 3 most popular primary languages were Python (n = 585), R (n = 118), and C++ (n = 88; Supplementary Table ST22). The 3 most frequently used languages with at least 1 file within a given repository were Python (n = 978), Shell (n = 712), and Dockerfile ( n = 350; Supplementary Table ST23). The 3 primary languages with the highest mean RQI were Go (n = 12, RQI = 3.15), Rust (n = 46, RQI = 3.06), and C (n = 42, RQI = 3.04; Supplementary Table ST24, Supplementary Fig. S15). The CI coverage for GO, Rust, and C was 91.7%, 84.8%, and 50% and their testing coverage was 100%, 87%, and 81%, respectively (Supplementary Table ST24). Where a comparison was possible, mean RQI was higher for repositories with CI than for those without CI in every language except GO, Typescript, and TeX (Supplementary Table ST25, Supplementary Fig. S16). Mean RQI was higher for repositories with tests in every language except for Shell, C++, and Julia (Supplementary Table ST25, Supplementary Fig. S17). Higher RQI scores for projects with CI and testing enabled suggest that these practices may contribute to the quality of software across different development frameworks.

### Repository Quality and Project Lifespan

To examine whether CI and testing help repositories maintain quality over time, repositories were binned into age cohorts, and mean RQI was compared across groups split by CI (Fig. 3A) and testing status (Fig. 3B). Repositories with CI maintained significantly higher RQI than those across all age cohorts (all cohorts p<0.0001), with the quality gap widening substantially as repositories aged (0.39 RQI 0-2 years to 0.75 RQI 10+ years; Supplementary Table ST26). A similar but less dramatic pattern was found for testing. The gaps between repositories with and without tests were significant across all cohorts (p<0.0001) but did not increase over time as much as CI (0.22 RQI 0-2 years to 0.38 RQI 10+ years; Supplementary Table ST27). The consistently larger and widening gap for CI suggests that the intentional implementation of CI can protect aging software from depreciation and aid in the long-term software support burden.

**Fig 3.**
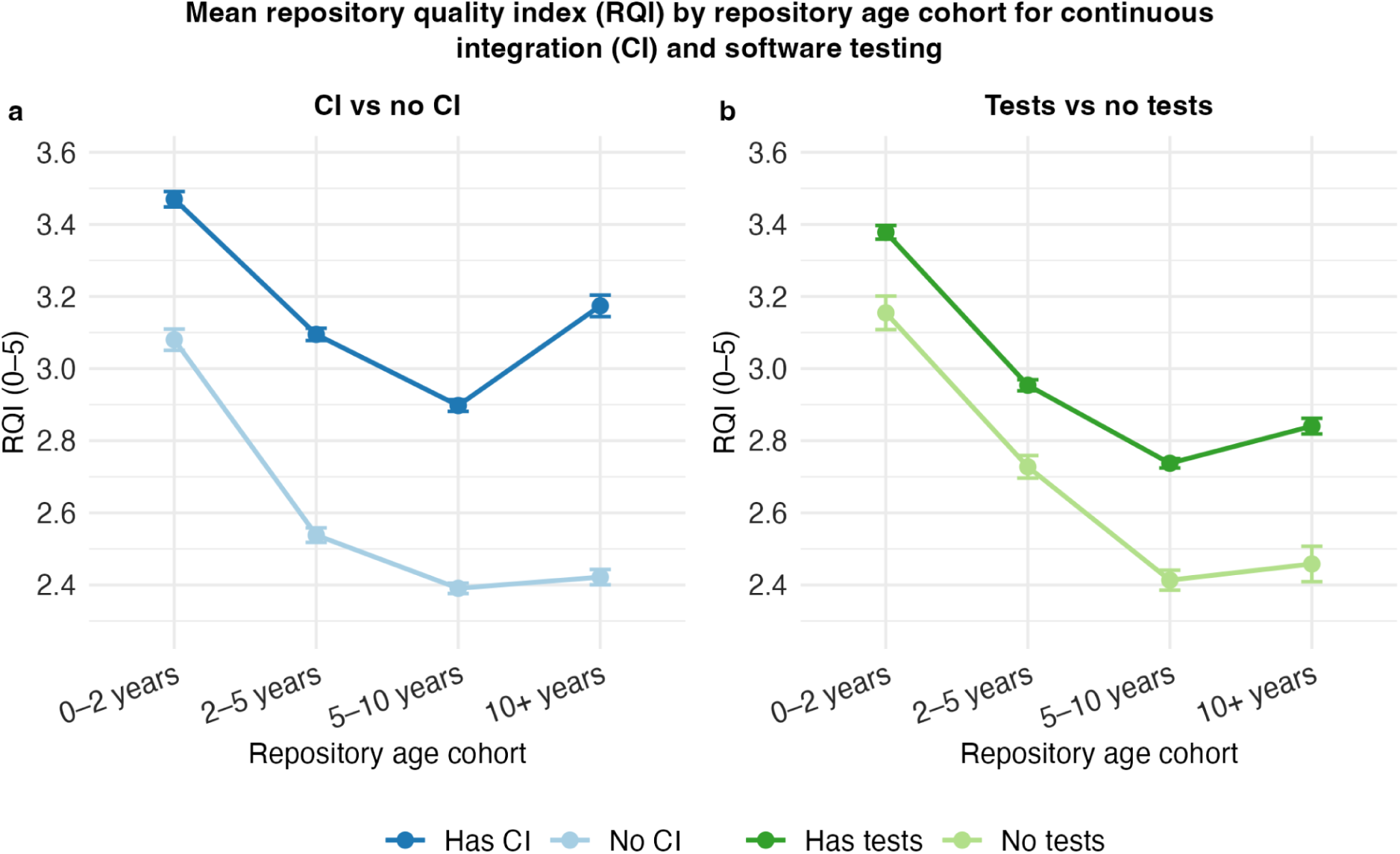
Mean RQI across Repository Age Cohorts by CI and Testing Status. (**a**) RQI for CI vs no CI. (**b**) RQI for test vs. no tests. Age is calculated as the years between the date of initial commit and the time of data collection.

### Software Quality as a Predictor of Publication and Funding

The relationship between RQI, publication, and funding outcomes was analyzed using projects from the bioinformatics group, where becoming a citable resource is likely a primary goal. Publication status was determined from the WoS hybrid search step in the data processing pipeline, where projects were marked as published if a citation was found and non-published if the search was unsuccessful across all search layers. This resulted in 1,450 projects, with 717 published and 733 unpublished projects (Fig. 4). A Wilcoxon test was then conducted on the RQI of the two groups, revealing a significant difference in the median RQI of published (2.82) and unpublished (2.49) projects (Fig. 4a). Funding status was determined from the OpenAlex and NIH RePORTER hybrid search step. This resulted in 711 bioinformatics projects with valid grant funding status indicators, with 473 being funded and 238 not funded (Fig. 4b). A Wilcoxon rank-sum test revealed a significant difference in median RQI between grant-funded (2.91) and not grant-funded (2.71) projects (p = 0.0413; Supplementary Table ST30). A Wilcoxon rank-sum test for all domains, which consisted of 674 (RQI Median = 2.96) grant-funded and 583 (RQI Median = 2.94) not grant-funded projects, was not significant (p = 0.559; Fig. S18, Table ST28-ST29). These results suggest that investing in RQI indicators can increase the likelihood of publication, which, for most scientific developers, is a central goal.

**Fig 4.**
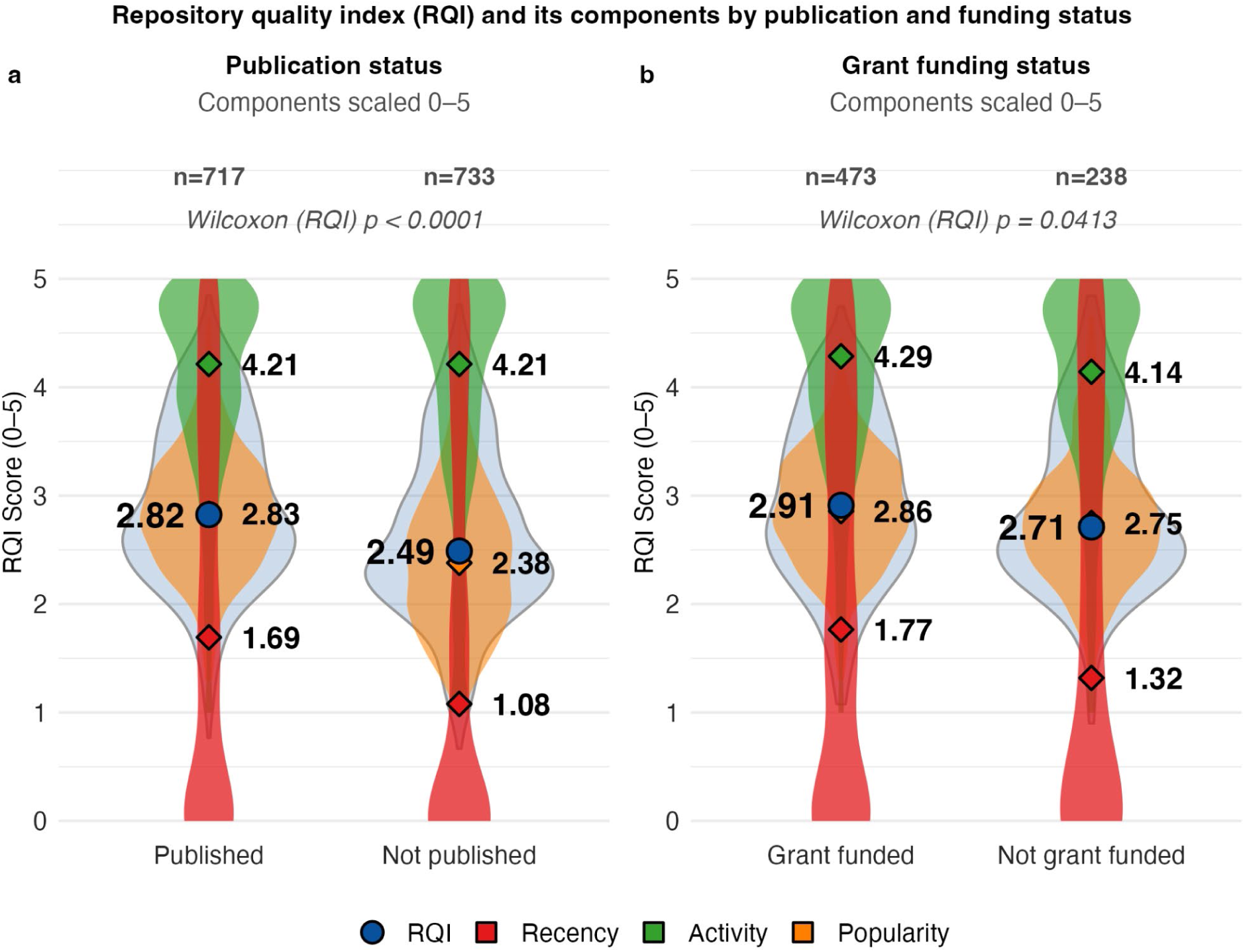
Median RQI by Publication and Funding Status for Bioinformatics Repositories. (**a**) Publication status for n=1,450 bioinformatics repositories. (**b**) Funding status for 711 bioinformatics repositories. The blue violin and circle represent the RQI distribution and median, respectively. Orange, green, and red violins show normalized Popularity, Activity, and Recency distributions, with diamonds marking their medians.

### Software Quality and Academic Citation Count

Higher repository quality was associated with greater academic citation count (Fig. 5). Repositories in the 1–2 and 2–2.5 RQI bins had median citation counts of 27.0 and 43.0, respectively, while higher-quality bins of 2.5–3.5, 3.5–4.5, and 4.5–5.0 had median counts of 39, 63, and 137. The overall difference between low (RQI ≤ 2.5, median = 27) and high (RQI > 2.5, median = 43) quality groups were statistically significant by the Wilcoxon rank-sum test (p = 0.003). Pairwise Wilcoxon tests revealed significant differences between most RQI bins. The 0-2 bin differed significantly from the 2-2.5 bin (p=0.015), the 2.5-3.5 bin (p=0.001), the 3.5-4.5 bin (p=0.003), and the 4.5-5.0 bin (p<0.001). The 2-2.5 bin also differed significantly from the 4.5-5 bin (p=0.003). Within the higher-quality group, the 2.5-3.5 bin differed from the 4.5-5 group (p=0.01), and the 3.5-4.5 bin was significantly different than the 4.5-5 bin (p=0.27). Log transformation was deemed to be the best fitting transformation for citation count (Supplementary Fig. S19, Supplementary Table ST31). Linear regression on log-transformed citation counts among published repositories confirmed this positive relationship (slope = 0.31, R² = 0.02, p < 0.001; Supplementary Fig. S20, Supplementary Tables ST32-ST33).

**Fig 5.**
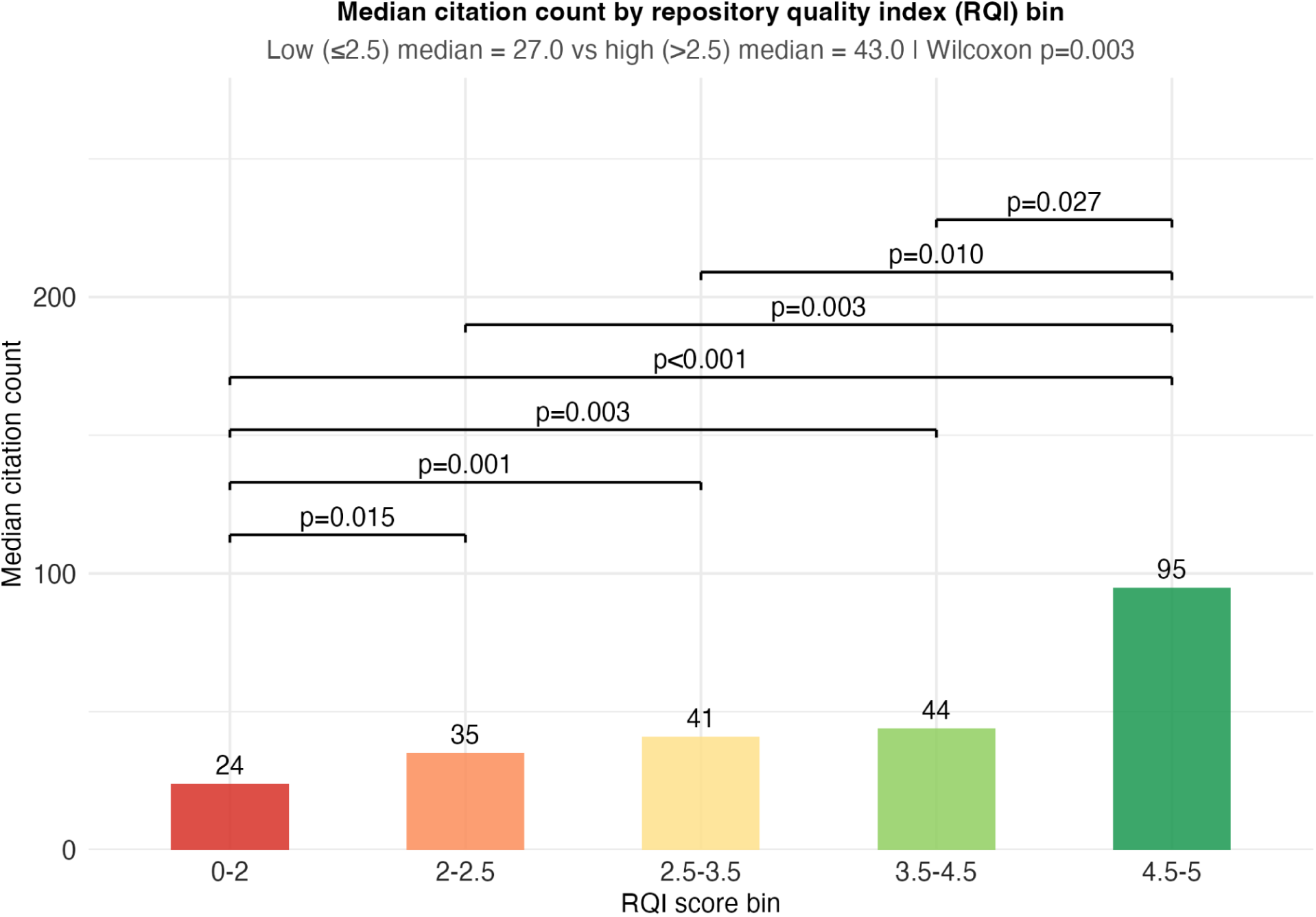
Median Citation Count by RQI Bin, Split by High/Low-Quality Groups. Wilcoxon Tests were performed between each bin and each quality group. Significant differences were found in the median RQI for published and unpublished projects.

### Resources and Data for Developers and Users

To provide an indicator for repository health, we developed a Repository Health Indicator Tool to help developers and users gauge how well project teams manage backlogs, recency, and community engagement. The web utility, developed in Python, uses the GitHub REST API to query data for a given GitHub repository. The user receives an RQI score for that repository, along with a breakdown of what drives the score. This tool was developed to help developers understand the strengths and weaknesses of their repositories, so they can identify which dimensions of their projects need attention. End-users can also employ this tool to assess whether a project is well-supported or on the brink of deprecation. By making it easy to understand the overall health of a software project, the tool helps users avoid wasting time and resources on tools that are no longer maintained. The utility also generates a markdown badge with a repository’s RQI score that can be added to a GitHub readme file. The tool is freely available at https://rqi.jakobilab.org.

## Discussion

The challenges facing the academic open-source community extend well beyond code quality and dependency management. The inherent design of the academic system could contribute to poor maintenance and development practices. A significant portion of this problem lies with grad student tenure. Early-career scientists, including PhD and MS students, are often assigned software development as a main project or task. The software is then created, published, and perhaps maintained for the remainder of the student’s time in that position. Once they move on, active maintenance often becomes less timely or ceases completely, since the very people best positioned to sustain it are usually not employed to maintain projects for extended periods of time. This is not a matter of individual neglect, but a reflection of how academic incentives are structured. Labs receive funding to build software, students graduate and move on, and once a project reaches publication, incentives to commit resources into project maintenance become limited. This leaves potentially crucial software to deteriorate as dependency chains break and computational environments evolve even without actively changing the codebase of the project itself. Pursuing new projects is, of course, necessary and commendable, but progress does not render prior work irrelevant. In reality, existing software plays a key role in the development of new methodologies, and allowing it to decay quietly undermines the very foundation that future research builds upon. Adopting industry-standard practices that are simple to implement would help prevent these important pieces of software from becoming legacy code. As we showed, in many cases, the implementation of these practices can even improve software as it ages past the initial development stage (Fig. 3).

The rise of AI-assisted development and prompt-driven development is lowering the barrier to entry for software creation, leading to an influx of software built and maintained by novices ^[12]^. Creating new novel software is especially enticing since there are significant publication and funding opportunities for quality software (Fig. 4). As these tools become ingrained in modern workflows, more individuals from non-computational backgrounds and limited formal engineering experience are likely to contribute to existing and new bioinformatics software. This increased accessibility has the potential to produce useful and novel tools but also risks introducing projects that lack foundational development practices, such as testing, CI, and security standards that traditionally trained developers would apply naturally. Ensuring that practices like CI are introduced early in the development process can serve as a guide for new developers and contributors by helping them ensure that their software is functional during development and sustainable beyond its initial release. The integration of AI into development workflows can also help close this gap by assisting in the creation of testing frameworks ^[13]^. These tools can give novice developers the guidance needed to write rigorous and effective tests, helping ensure their software remains stable and maintainable over time. While AI is not a substitute for a dedicated testing engineer, it can meaningfully improve the outlook and performance of software potentially produced by small or inexperienced teams ^[14]^.

The reproducibility crisis in science extends beyond the wet lab and into the software that powers the dry lab ^[15]^. As digital ecosystems evolve, existing software is susceptible to unexpected changes in behavior. How projects address deviations from original expectations can vary widely. For example, a floating-point calculation differing by 10⁻^10^ might be inconsequential for one application yet produce misleading results in another. Testing and CI pipelines can help by running the software against expected outputs and alerting the development team when an element of the software is acting unexpectedly, allowing them to assess and address the change appropriately and even improve upon the software’s original formula (Fig. 3).

Another crux in the scientific software reproducibility crisis is the availability of code for both active and inactive projects. Making consistently tested code available to the public not only ensures that results are consistent but also gives developers the ability to confidently contribute to or utilize code that is stable. Instead of having to start from scratch, developers can build upon approaches that are already verified by up-to-par development practices. This not only ensures that software is reproducible but also ready for future iterations of improvement and innovation. Using central and open repositories for software development and deployment also helps to hold on to legacy code from historic software that otherwise might disappear when lab websites move or get deleted.

We found that popular software projects that invest resources into consistent and recent updates have better publication, citation, and grant outcomes than those with lower levels of software quality indicators (Figs. 4, 5). Our findings indicate that adopting practices directly correlated with software quality, such as CI, has the potential to positively impact the academic performance of software projects and the labs that create and maintain them. Monetary support, citations, and publications can be considered direct indicators of a lab’s success and health, and investments in high-quality software development practices lead to real benefits for the lab and the members developing the software. Conversely, software developed without these quality indicators in mind risks missing grants, publications, and citations, which are all relevant measures for most modern researchers and organizations. Utilizing available tools, such as GitHub’s actions suite (https://github.com/features/actions) or GitLab’s CI suite (https://docs.gitlab.com/ci/), can ensure that software projects can meet these standards that ultimately predict a lab’s status. Many of these tools are free or low-cost for open-source developers and academic institutions.

The findings of our study suggest that academic software development might benefit significantly from the adoption of industry-standard practices of robust testing and CI. Projects that incorporate CI demonstrate improvements in both software quality and project longevity (Fig. 2). The value of integrative testing extends beyond these long-term measures. Effectively implemented CI pipelines can detect breaking points within multiple functional areas of a codebase spanning diverse computational environments. Consequently, a large portion of the responsibility of detecting broken dependency chains, faulty logic, and defective code shifts from the user community (reactive handling) to an automated process operating within controlled environments (proactive testing) that developers can directly investigate. Overall, this can help to deliver code with fewer issues, thus making long-term maintenance more manageable.

## Conclusion

Utilizing development tools that uphold the stability and support of software ensure that the effort, time, and resources that go into creating them are worthwhile. Our findings show that intentionally implemented workflows for CI profoundly impact the lifespan of software and the academic and financial success of the organizations maintaining them. CI, in this case, represents an overall consideration for the whole software lifecycle, specifically the support, maintenance, and end-of-life stages. Alongside CI, we believe there are many other approaches to long-term support that developers should consider. The responsibility throughout the software lifecycle falls heavily on the scientific developer, since scientific discovery and progress are dependent upon the stability and accuracy of the software. These development tools share that responsibility, keeping maintenance and further development manageable. In some cases, that support even improves the software over time. Embracing these practices not only enhances the software itself but also upholds the integrity and continuity of computational science and its community.

## Data Availability

We developed an R Shiny exploration tool that lets users view and interact with the main and supplementary figures of this study. The software is available at https://shiny.jakobilab.org/rqi.

All data, figures, code, and the supplementary markdown file are available at https://GitHub.com/jakobilab/rqi-study. The supplementary data is also available in markdown format at https://jakobilab.org/rqi-study/.

Our Repository Quality Indicator is a web-based software that gauges relative backlog health, recency, and community engagement for a given public GitHub repository. It is available at https://rqi.jakobilab.org. All data and software is licensed under the GNU GPL v3 license.

## Author Contributions

W.J.C.M. led the manuscript preparation, methodology, and data analysis. M.S. and N.G. contributed to the writing and interpretation of data. T.J. was responsible for overall project conceptualization, supervision, bioinformatics direction, and manuscript refinement.

## Additional Information

### Competing Interests

The authors declare no competing interests.

## Notes

https://rqi.jakobilab.org

https://shiny.jakobilab.org/rqi

https://jakobilab.org/rqi-study

### Competing Interest Statement

The authors have declared no competing interest.

### Funding

- National Human Genome Research Institute: R01HG013124
- Arizona Biomedical Research Centre: RFGA2022-010-11

