## Supplementray File 1 for "Continuous Integration and Software Quality in Scientific Software: A Large-Scale Empirical Analysis of GitHub Repositories"

Online version available at jakobilab.org/rqi-study

Walker J. Compton Mellon, Makena Sy, Navya Gangrade, Tobias Jakobi

2026-09-30

#### Contents


**1. Data** **2**

**2. RQI Scoring** **3**

**3. Summary Statistics** **4**

**4. Analysis of Normality for RQI** **5**

**5. ANOVA and Tukey Tests** **8**

**6. Significant Comparisons** **9**

**7. Mean Scores by Type** **10**

**8. Group Means** **11**

**9. Radar Plots** **12**

**10. CI and Test Prevalence by Quality** **18**

**11. CI and Tests by Type** **21**

**12. Languages** **24**

**13. Repository Age** **37**

**14. Citations and RQI** **41**

**15. Publication Status** **42**

**16. Funding** **43**

**17. Citation Models** **46**

### 1. Data

**Table ST1. Repositories per domain, before and after filtering**

| Domain | Raw | After filtering |
|---|---|---|
| open_source | 162730 | 10864 |
| bioinformatics | 39539 | 1493 |
| astrophysics | 3254 | 243 |
| software_engineering | 95429 | 1088 |
| image_recognition | 39912 | 149 |

#### 2. RQI Scoring

RQI sub-scores use piecewise-linear interpolation between fixed anchor points, so scores do not depend on the sample.

$$\mathrm{RQI}_i = \frac{1}{3}\Big(f_{\mathrm{recency}}(\mathrm{days\_since\_last\_commit}_i) + f_{\mathrm{issues}}(\mathrm{median\_issue\_days\_open}_i) + f_{\mathrm{popularity}}\big(\log_{10}(1+\mathrm{stars}_i+\mathrm{forks}_i+\mathrm{watchers}_i)\big)\Big)$$

where each $f$ interpolates between fixed anchors on a 0–5 scale. Values beyond the first or last anchor take the score of that anchor.

**Table ST2. RQI scoring anchors.** Scores are linearly interpolated between anchors.

| Recency: days since last commit | 0 | 14 | 30 | 60 | 90 | 180 | 365 | 730 |
|---|---|---|---|---|---|---|---|---|
| $f_{\mathrm{recency}}$ | 5.0 | 4.5 | 4.0 | 3.5 | 3.0 | 2.0 | 1.3 | 0.0 |

| Backlog: median days issues stay open | 0 | 7 | 14 | 30 | 60 | 120 | 365 |
|---|---|---|---|---|---|---|---|
| $f_{\mathrm{issues}}$ | 5.0 | 4.5 | 4.0 | 3.5 | 3.0 | 2.0 | 1.0 |

| Popularity: $\log_{10}(1 + \mathrm{stars} + \mathrm{forks} + \mathrm{watchers})$ | 0 | 1 | 2 | 3 | 4 |
|---|---|---|---|---|---|
| $f_{\mathrm{popularity}}$ | 1.0 | 2.0 | 3.0 | 4.0 | 5.0 |

Missing values: a repository with no issue-closure median is scored as 365 days open (backlog 1.0).

### 3. Summary Statistics

#### 3.1 Overall and by Type

**Table ST3. Overall repository summary**

| Statistic | Value |
|---|---|
| Repositories | 13,670 |
| Without CI | 5,085 |
| Without tests | 1,742 |
| CI coverage (%) | 62.8 |
| Test coverage (%) | 87.3 |
| Neither CI nor tests (%) | 7.5 |
| Mean RQI | 2.88 |
| SD of RQI | 0.90 |
| Minimum RQI | 0.67 |
| Maximum RQI | 5.00 |

**Table ST4. Summary by repository type**

| Type | n | CI (%) | Tests (%) | Neither (%) | Mean RQI | SD RQI |
|---|---|---|---|---|---|---|
| astrophysics | 238 | 73.9 | 92.4 | 4.6 | 2.97 | 0.84 |
| open_source | 10773 | 64.4 | 86.8 | 7.3 | 2.96 | 0.92 |
| bioinformatics | 1456 | 65.9 | 89.3 | 7.7 | 2.79 | 0.78 |
| image_recognition | 144 | 36.1 | 77.8 | 18.1 | 2.56 | 0.74 |
| software_engineering | 1059 | 43.6 | 89.0 | 8.6 | 2.22 | 0.61 |

### 4. Analysis of Normality for RQI

#### 4.1 RQI by Repository Type

**Fig S1. RQI distribution by repository type.** Stacked histogram with density overlays.

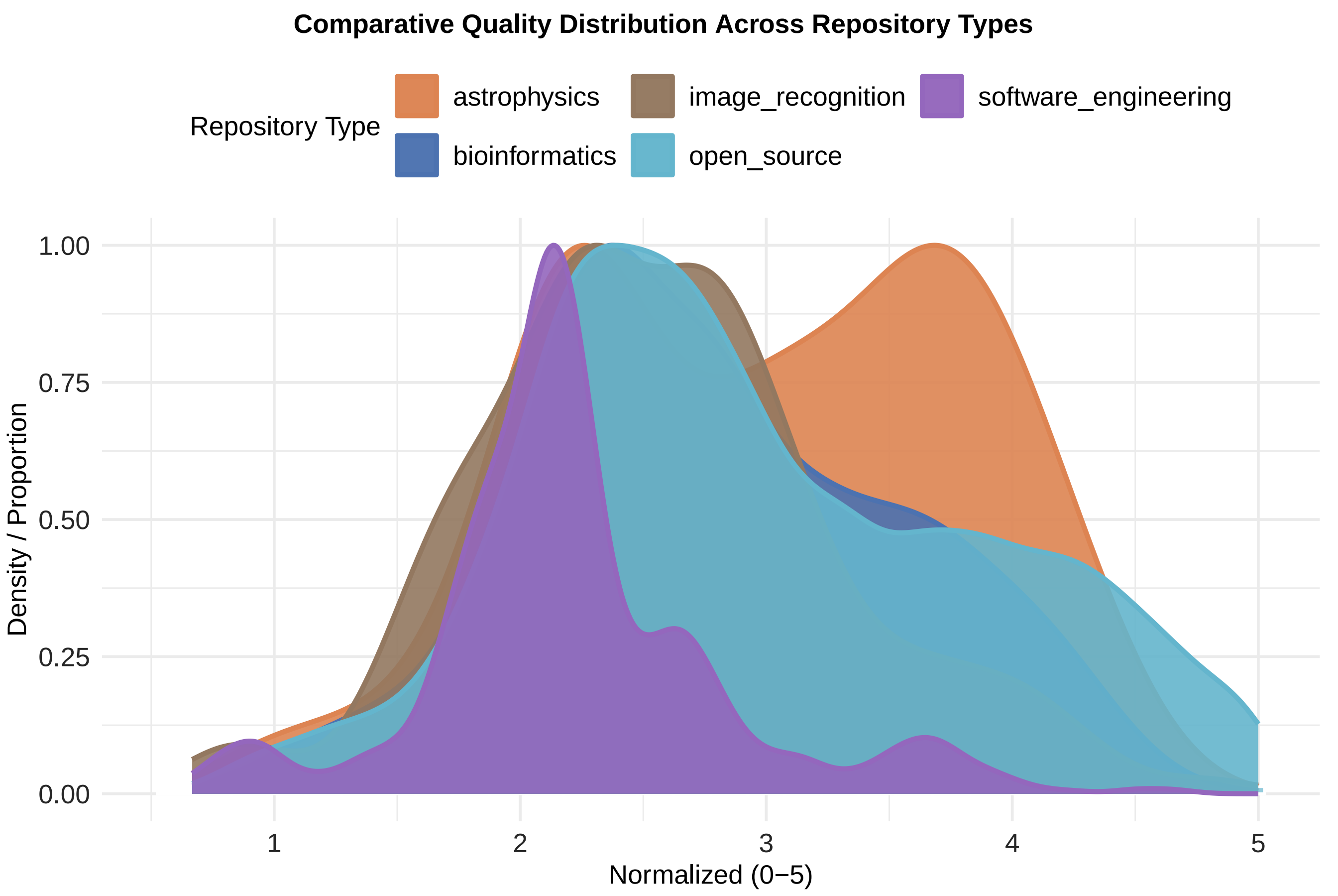

##### 4.2 Overall RQI Distribution

**Fig S2. Overall RQI distribution and Q–Q plot.** Dashed line marks the mean.

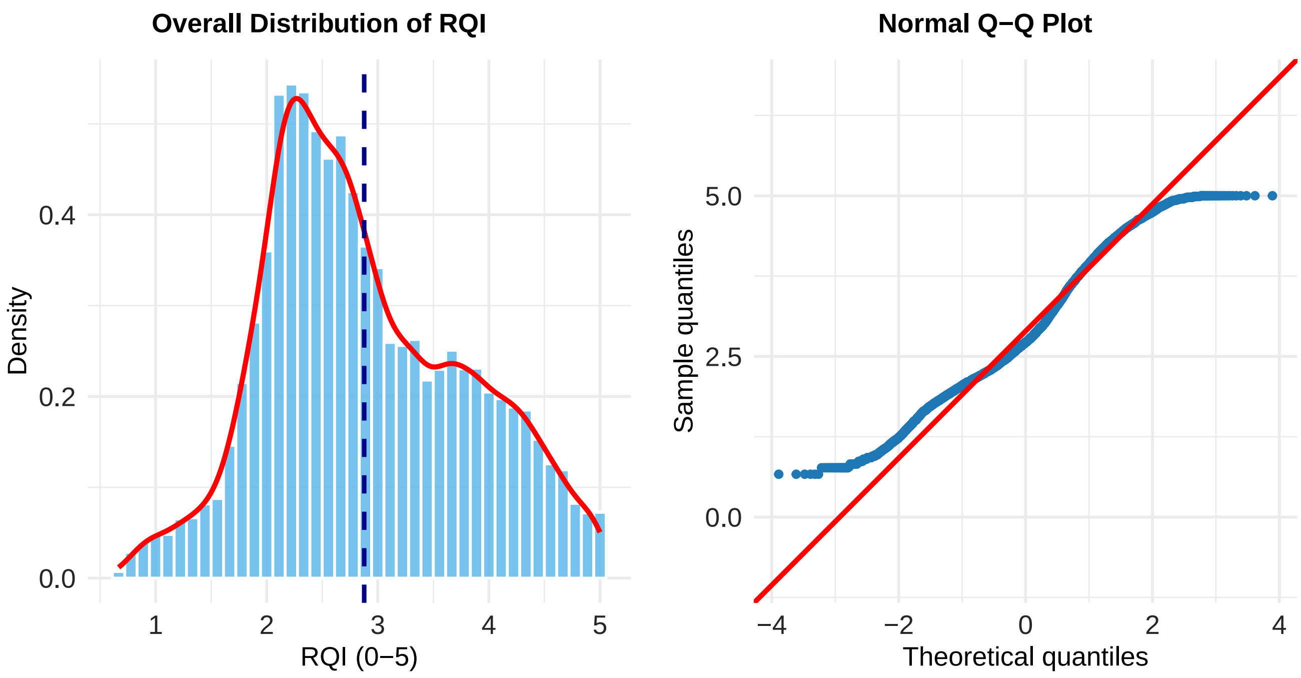


**Table ST5. Overall RQI distribution: summary statistics and normality diagnostics**

| n | Min | Q1 | Median | Mean | Q3 | Max | SD | Skewness | KS D | KS p |
|---|---|---|---|---|---|---|---|---|---|---|
| 13670 | 0.667 | 2.217 | 2.715 | 2.878 | 3.55 | 5 | 0.904 | 0.338 | 0.073 | <0.001 |

##### 4.3 RQI Distribution by Type

**Fig S3. RQI distribution by repository type.** Kolmogorov–Smirnov statistics and skewness shown per type.

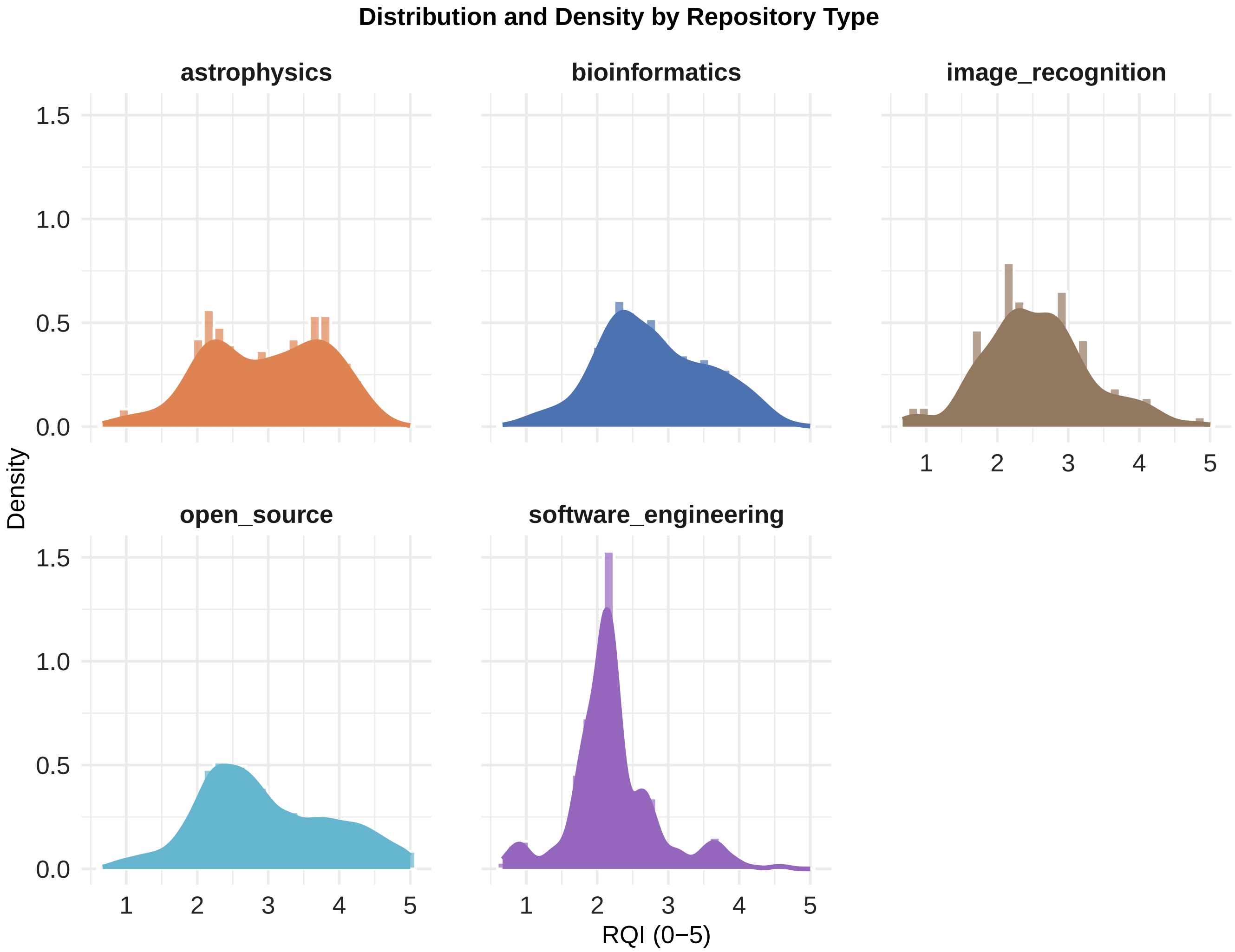


**Table ST6. Normality diagnostics for RQI by repository type**

| Type | n | KS D | KS p | Skewness |
|---|---|---|---|---|
| astrophysics | 238 | 0.082 | 0.084 | -0.191 |
| bioinformatics | 1456 | 0.055 | $<$0.001 | 0.170 |
| image_recognition | 144 | 0.064 | 0.603 | 0.311 |
| open_source | 10773 | 0.071 | $<$0.001 | 0.278 |
| software_engineering | 1059 | 0.141 | $<$0.001 | 0.677 |

P values indicate non-normality, but the plots and low D values suggest normality is adequate for ANOVA.

### 5. ANOVA and Tukey Tests

**Table ST7. One-way ANOVA of RQI by repository type**

| Term | Df | Sum Sq | Mean Sq | F | p |
|---|---|---|---|---|---|
| Repository type | 4 | 555.98 | 138.99 | 178.97 | <0.001 |
| Residuals | 13665 | 10612.86 | 0.78 | | |

**Table ST8. Tukey HSD pairwise comparisons**

| Comparison | Difference | CI lower | CI upper | Adj. p |
|---|---|---|---|---|
| bioinformatics vs. astrophysics | -0.185 | -0.353 | -0.017 | 0.022 |
| image_recognition vs. astrophysics | -0.409 | -0.663 | -0.155 | <0.001 |
| open_source vs. astrophysics | -0.017 | -0.175 | 0.140 | 0.998 |
| software_engineering vs. astrophysics | -0.756 | -0.928 | -0.583 | <0.001 |
| image_recognition vs. bioinformatics | -0.224 | -0.434 | -0.014 | 0.030 |
| open_source vs. bioinformatics | 0.168 | 0.101 | 0.235 | <0.001 |
| software_engineering vs. bioinformatics | -0.571 | -0.668 | -0.474 | <0.001 |
| open_source vs. image_recognition | 0.392 | 0.190 | 0.594 | <0.001 |
| software_engineering vs. image_recognition | -0.347 | -0.560 | -0.133 | <0.001 |
| software_engineering vs. open_source | -0.739 | -0.816 | -0.661 | <0.001 |

### 6. Significant Comparisons

**Table ST9. Tukey HSD comparisons with adjusted p < 0.05**

| Group1 | Group2 | Difference | CI lower | CI upper | Adj. p |
|---|---|---|---|---|---|
| bioinformatics | astrophysics | -0.185 | -0.353 | -0.017 | 0.022 |
| image_recognition | astrophysics | -0.409 | -0.663 | -0.155 | <0.001 |
| software_engineering | astrophysics | -0.756 | -0.928 | -0.583 | <0.001 |
| image_recognition | bioinformatics | -0.224 | -0.434 | -0.014 | 0.030 |
| open_source | bioinformatics | 0.168 | 0.101 | 0.235 | <0.001 |
| software_engineering | bioinformatics | -0.571 | -0.668 | -0.474 | <0.001 |
| open_source | image_recognition | 0.392 | 0.190 | 0.594 | <0.001 |
| software_engineering | image_recognition | -0.347 | -0.560 | -0.133 | <0.001 |
| software_engineering | open_source | -0.739 | -0.816 | -0.661 | <0.001 |

### 7. Mean Scores by Type

**Table 1 (main text). Mean scores by repository type.**

| Type | Mean RQI | Mean Recency | Mean Backlog | Mean Popularity |
|---|---|---|---|---|
| astrophysics | 2.97 | 92.09 | 0.21 | 3.39 |
| bioinformatics | 2.79 | 88.10 | 0.21 | 3.74 |
| image_recognition | 2.56 | 78.86 | 0.32 | 3.99 |
| open_source | 2.96 | 86.31 | 0.29 | 4.61 |
| software_engineering | 2.22 | 78.28 | 0.33 | 1.55 |

### 8. Group Means

**Fig S4. Mean RQI, bioinformatics vs other domains.** Means with 95% CIs; brackets give Tukey-adjusted p values for significant comparisons.

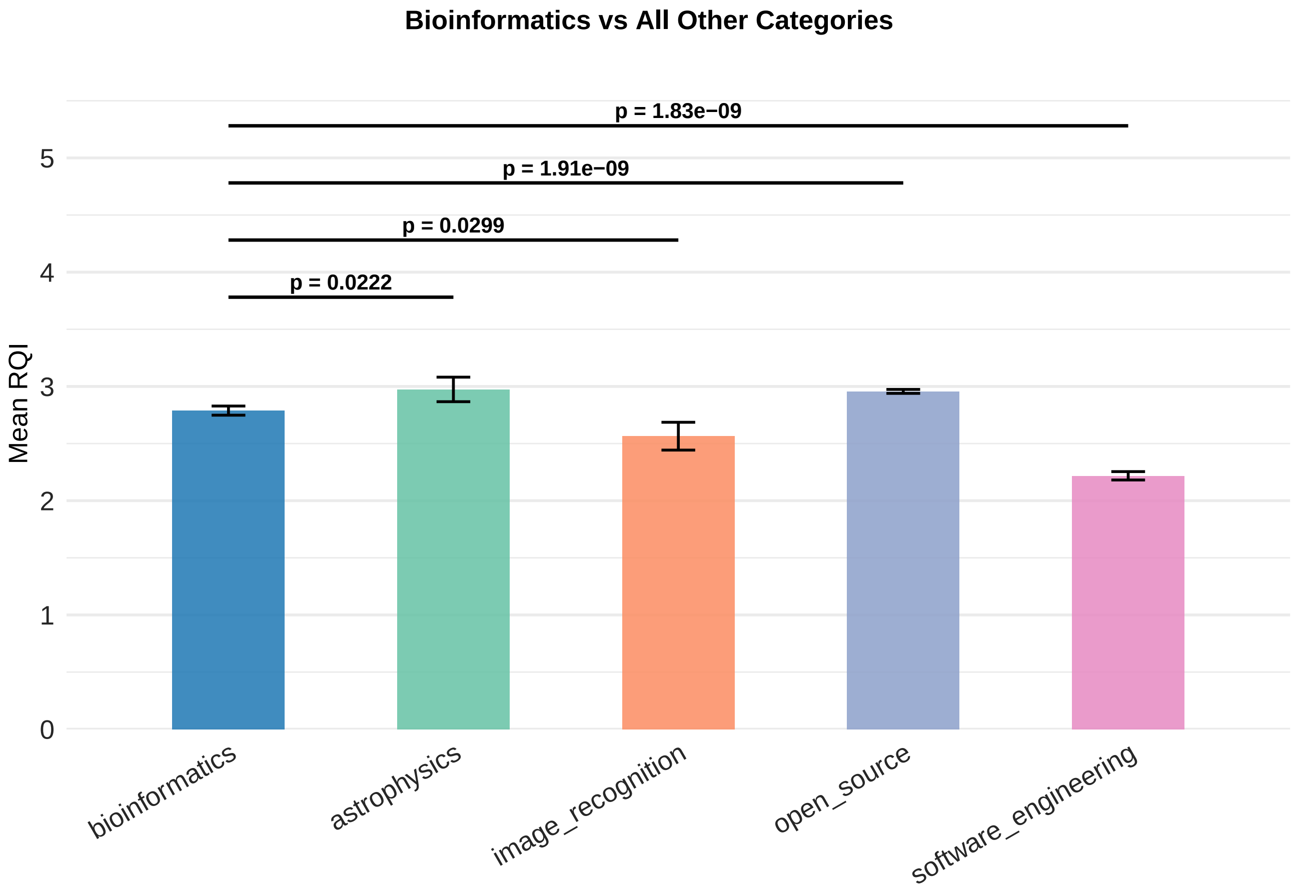

### 9. Radar Plots

#### 9.1 Top, Median, Bottom 100

**Fig 1 (main text). Radar plots of the top, median, and bottom 100 repositories by RQI.** Panels a–c; values scaled 0–100 (100 = best).

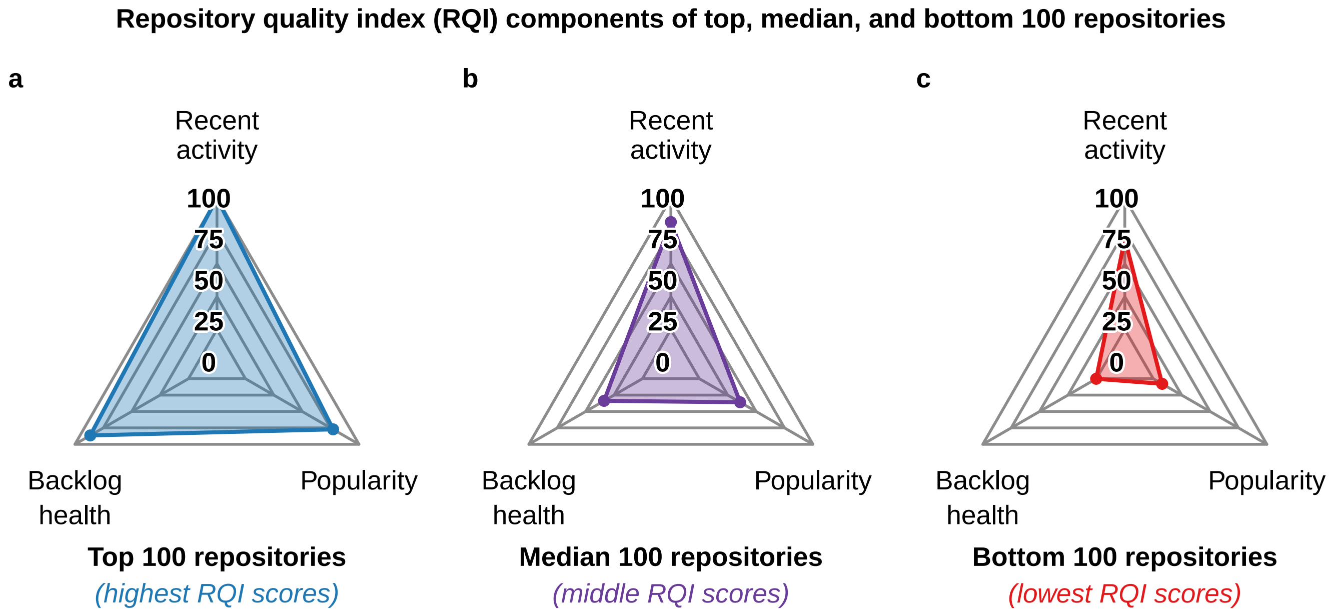


**Table ST10.** Fig 1 values (0–100).

Table 2: Radar plot values for Fig 1 (0-100 scale; outer edge = 100 = best).

| Group | Recent Activity | Backlog Health | Popularity |
|---|---|---|---|
| Top 100 (highest RQI) | 100.0 | 86.5 | 77.2 |
| Median 100 | 81.8 | 33.8 | 36.0 |
| Bottom 100 (lowest RQI) | 69.0 | 0.2 | 7.9 |

#### 9.2 Bioinformatics Top, Median, Bottom 100

**Fig S5. Radar plots of the top, median, and bottom 100 bioinformatics repositories.**

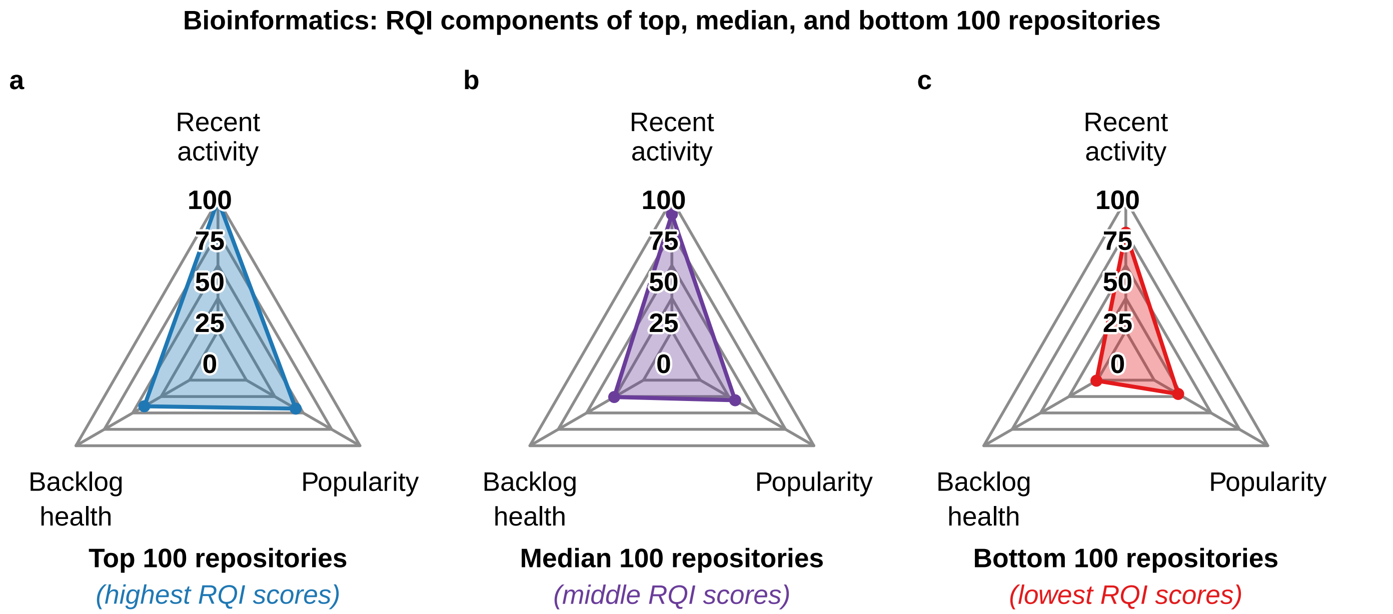


**Table ST11.** Fig S5 values (0–100).

Table 3: Radar plot values for Fig S5 (bioinformatics, 0-100 scale).

| Group | Recent Activity | Backlog Health | Popularity |
|---|---|---|---|
| Top 100 (highest RQI) | 99.8 | 39.8 | 43.4 |
| Median 100 | 89.0 | 25.7 | 30.7 |
| Bottom 100 (lowest RQI) | 74.9 | 0.8 | 21.1 |

#### 9.3 Profile by Domain

**Fig S6. Average RQI component profile by domain.**

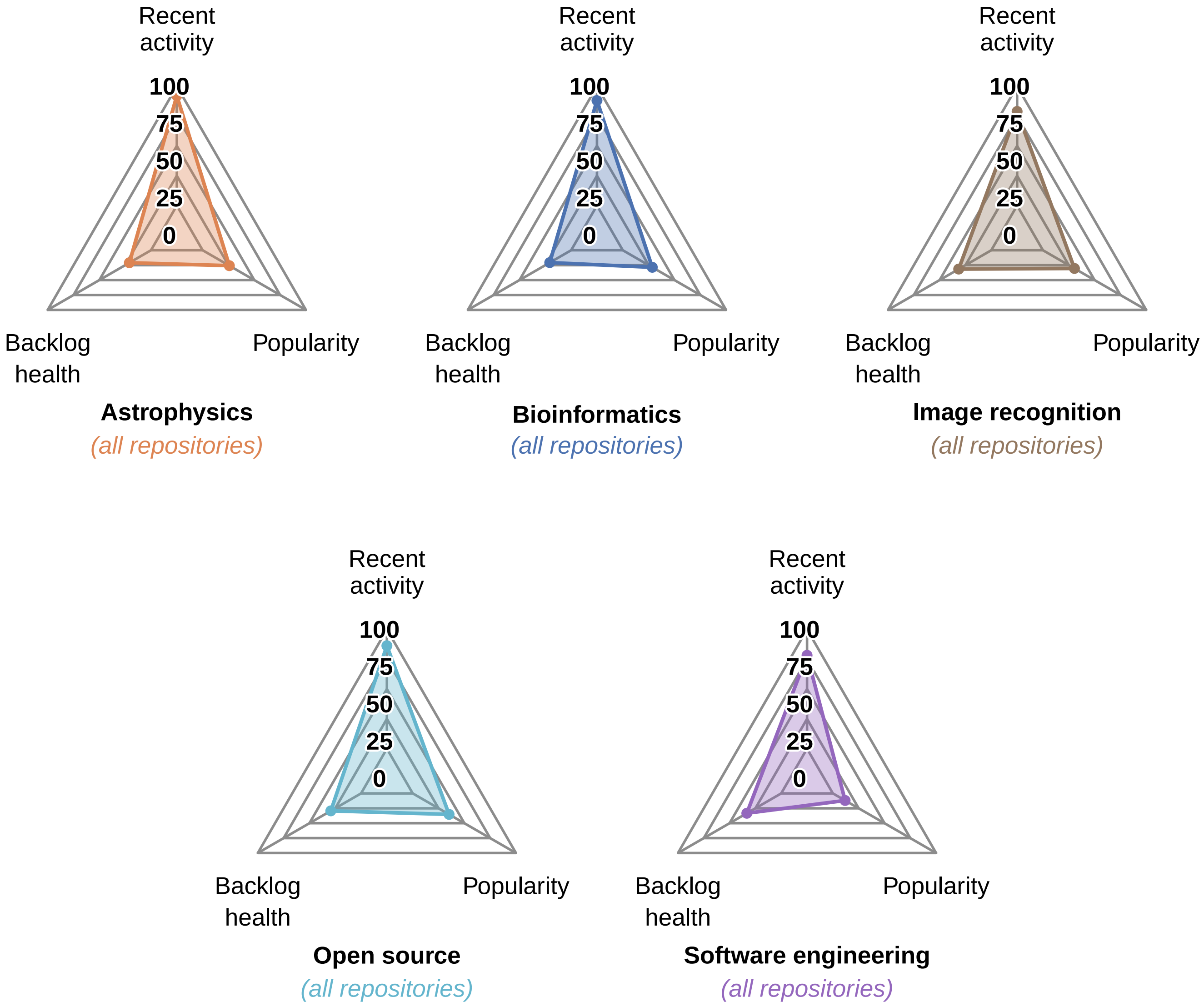


**Table ST12.** Fig S6 values (0–100).

Table 4: Radar plot values for Fig S6, by domain (0-100 scale).

| Domain | Recent Activity | Backlog Health | Popularity |
|---|---|---|---|
| astrophysics | 92.1 | 20.9 | 25.9 |
| bioinformatics | 88.1 | 20.7 | 28.6 |
| image_recognition | 78.9 | 31.6 | 30.5 |
| open_source | 86.3 | 29.3 | 35.2 |
| software_engineering | 78.3 | 33.3 | 11.9 |

#### 9.4 Top vs Bottom 20 by Domain

**Fig S7. Top 20 vs bottom 20 repositories by domain.** Left: highest RQI; right: lowest.

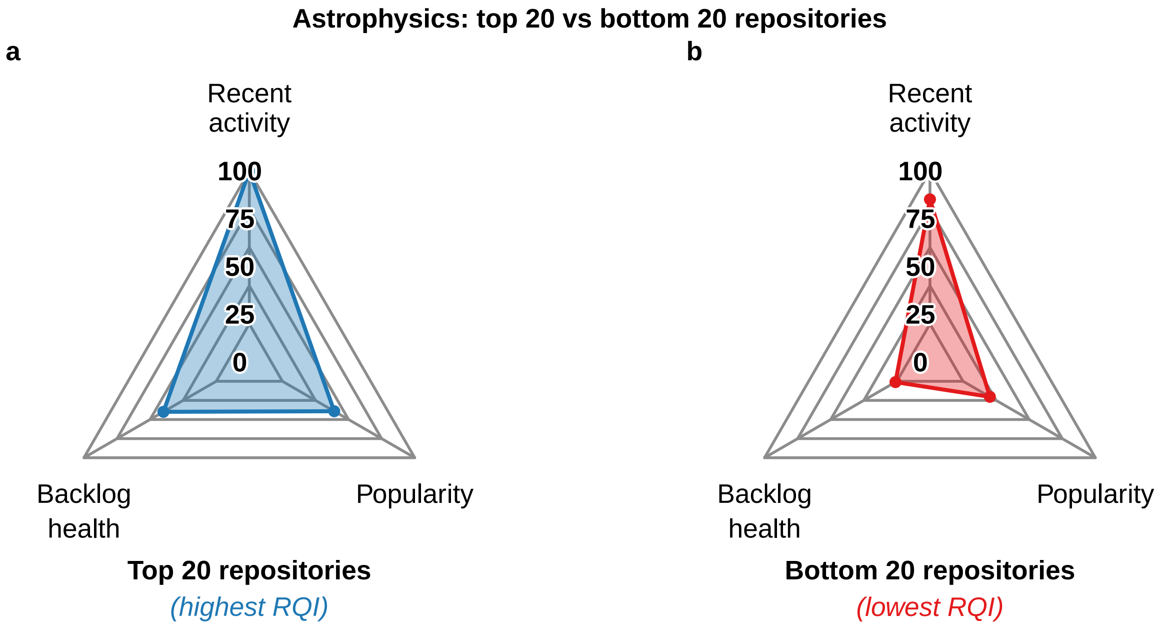


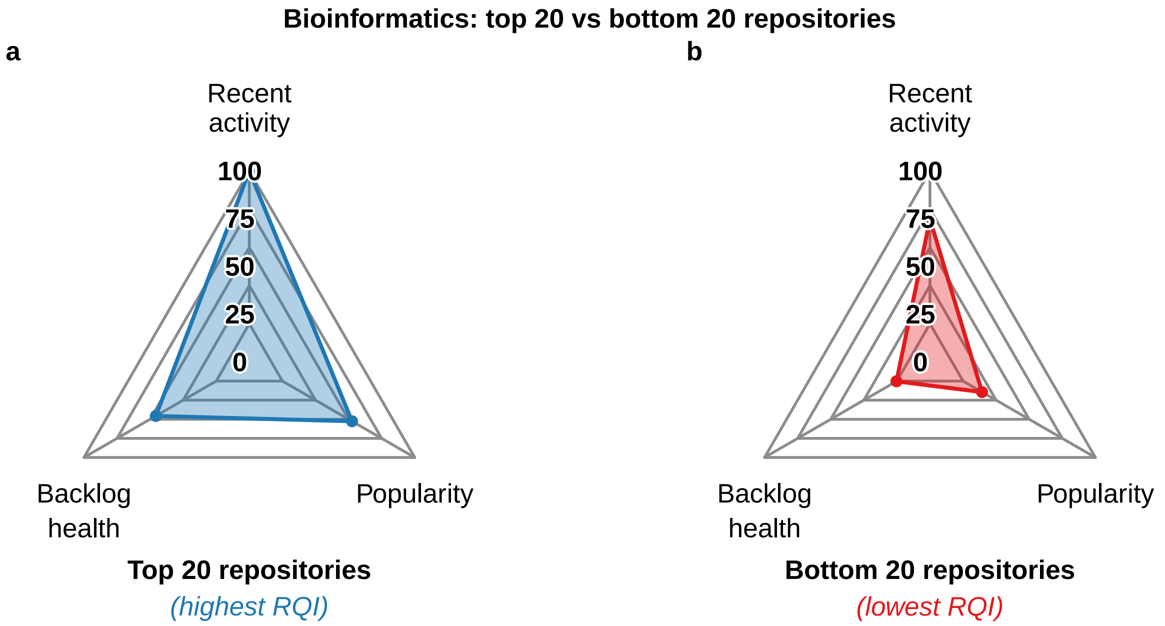

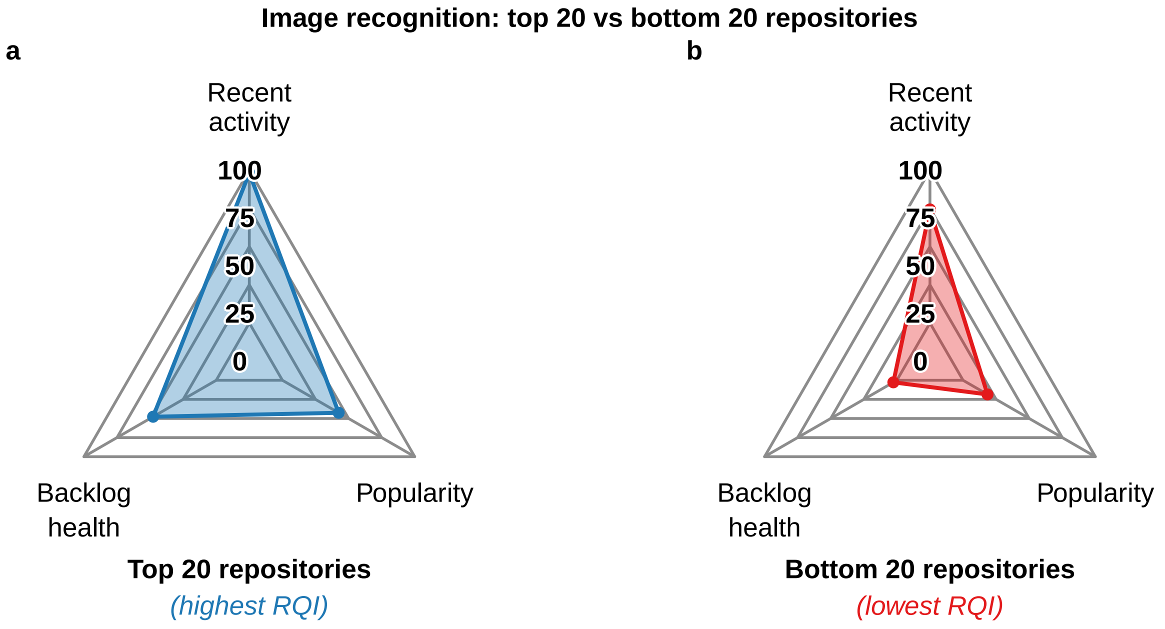
Image recognition: top 20 vs bottom 20 repositories
a
b
Recent activity
100
75
50
25
0
Backlog health
Popularity
Top 20 repositories
(highest RQI)
Recent activity
100
75
50
25
0
Backlog health
Popularity
Bottom 20 repositories
(lowest RQI)

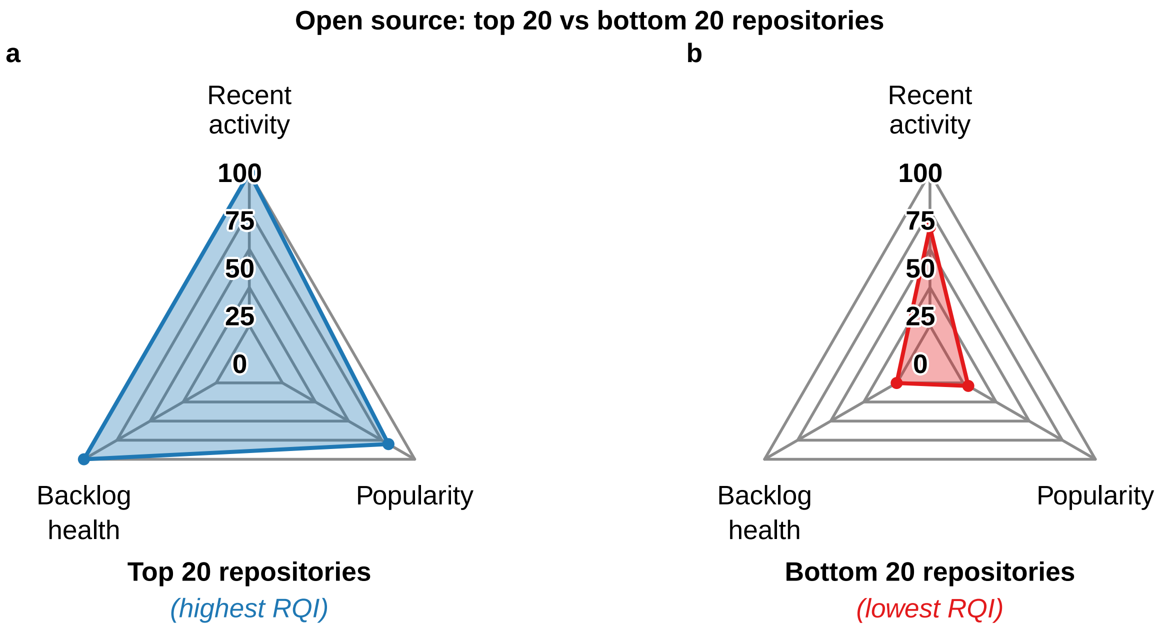
Open source: top 20 vs bottom 20 repositories
a
b
Recent activity
100
75
50
25
0
Backlog health
Popularity
Top 20 repositories
(highest RQI)
Recent activity
100
75
50
25
0
Backlog health
Popularity
Bottom 20 repositories
(lowest RQI)

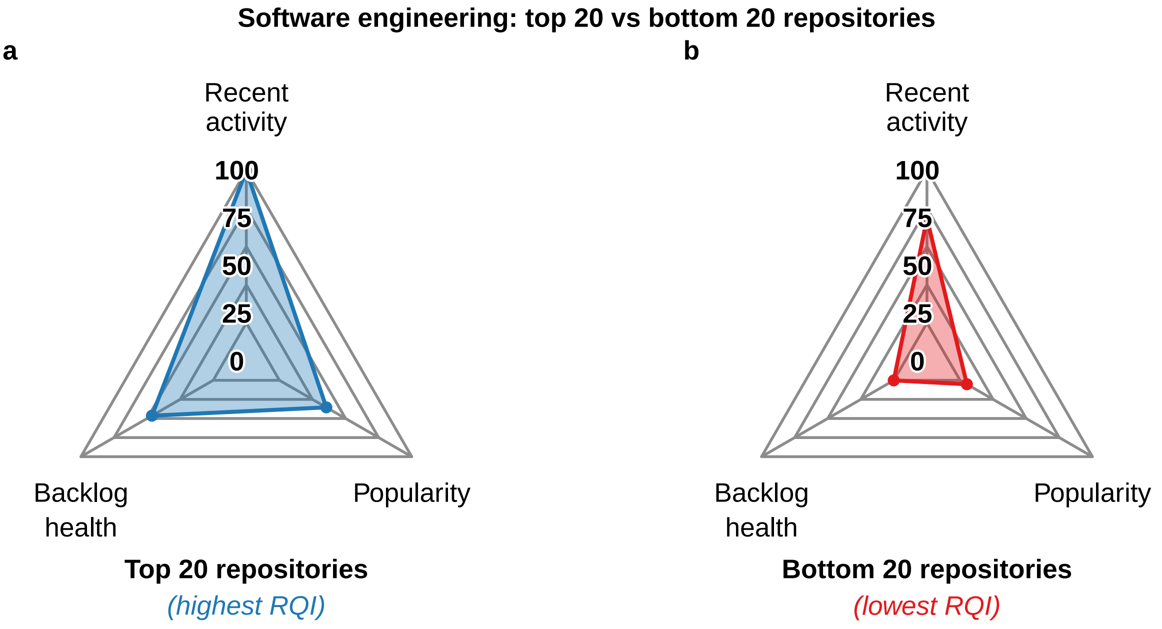


**Table ST13.** Fig S7 values (0–100).

Table 5: Radar plot values for Fig S7, top 20 vs bottom 20 by domain (0-100 scale).

| Domain | Group | Recent Activity | Backlog Health | Popularity |
|---|---|---|---|---|
| astrophysics | Top 20 | 99.9 | 39.9 | 39.2 |
| astrophysics | Bottom 20 | 81.5 | 1.1 | 20.3 |
| bioinformatics | Top 20 | 100.0 | 45.6 | 52.6 |
| bioinformatics | Bottom 20 | 68.4 | 0.3 | 14.4 |
| image_recognition | Top 20 | 98.2 | 47.7 | 42.5 |
| image_recognition | Bottom 20 | 74.3 | 2.6 | 18.5 |
| open_source | Top 20 | 100.0 | 100.0 | 80.2 |
| open_source | Bottom 20 | 64.4 | 0.2 | 4.0 |
| software_engineering | Top 20 | 99.7 | 46.3 | 35.5 |
| software_engineering | Bottom 20 | 68.4 | 0.1 | 5.2 |

### 10. CI and Test Prevalence by Quality

#### 10.1 CI and Tests by RQI

**Fig 2 (main text). CI and testing prevalence by RQI.** Bins of 0.15 RQI (n 5); point size shows bin n; lines are OLS fits.

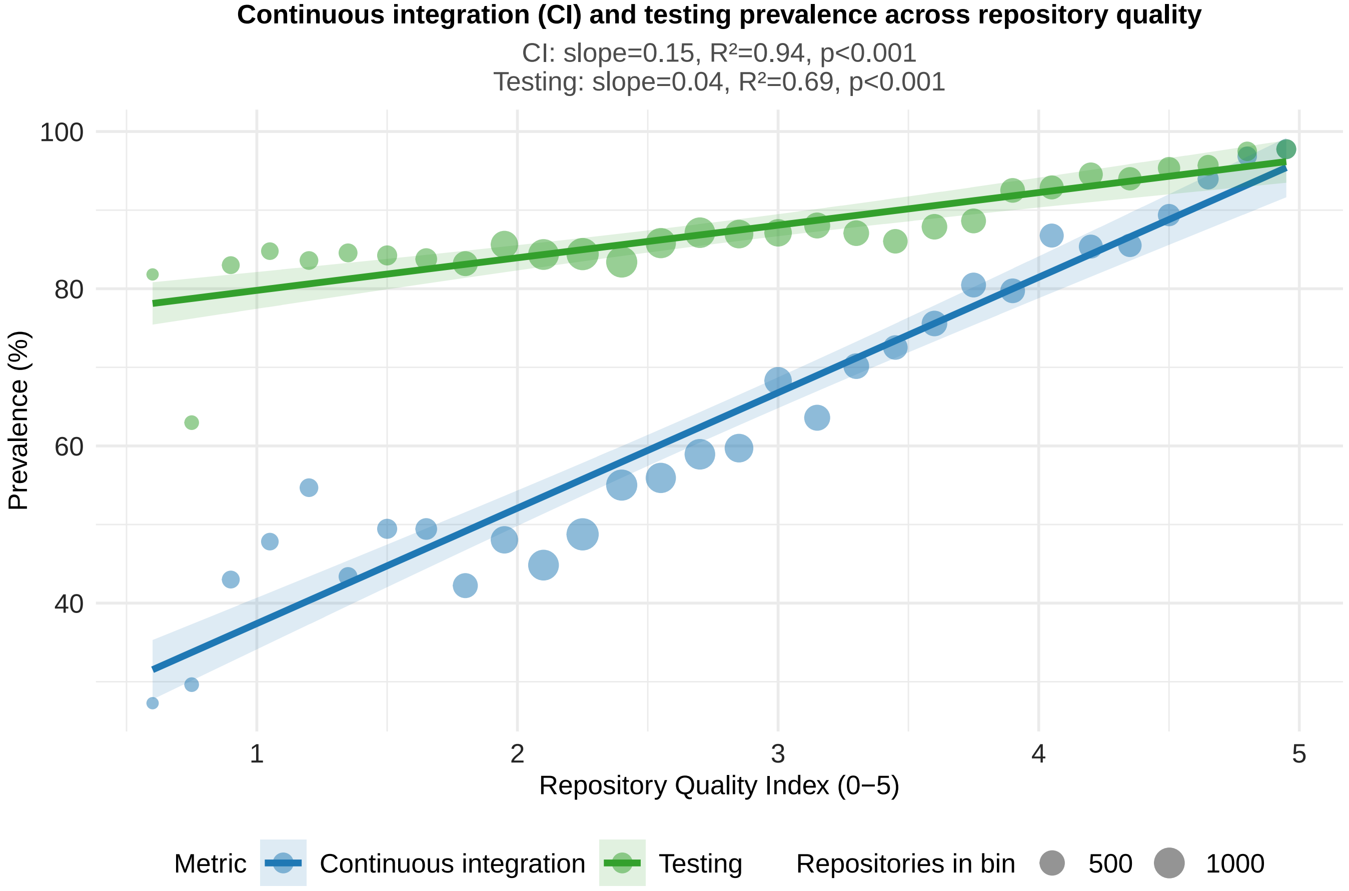

#### 10.2 Repos per RQI Bin

**Table ST14. Repositories per RQI bin (bin width 0.15)**

| RQI bin | Repositories |
|---|---|
| 0.60 | 11 |
| 0.75 | 27 |
| 0.90 | 100 |
| 1.05 | 92 |
| 1.20 | 128 |
| 1.35 | 136 |
| 1.50 | 184 |
| 1.65 | 265 |
| 1.80 | 476 |
| 1.95 | 668 |
| 2.10 | 997 |
| 2.25 | 1161 |
| 2.40 | 1036 |
| 2.55 | 944 |
| 2.70 | 979 |
| 2.85 | 782 |
| 3.00 | 675 |
| 3.15 | 552 |
| 3.30 | 526 |
| 3.45 | 444 |
| 3.60 | 520 |
| 3.75 | 466 |
| 3.90 | 454 |
| 4.05 | 408 |
| 4.20 | 403 |
| 4.35 | 366 |
| 4.50 | 301 |
| 4.65 | 232 |
| 4.80 | 159 |
| 4.95 | 178 |

#### 10.3 CI and Tests by Quality Tier

**Fig S8. CI and testing prevalence by RQI tier.** Low (RQI 2), mid, and high (RQI 4) tiers; Holm-adjusted pairwise tests shown.

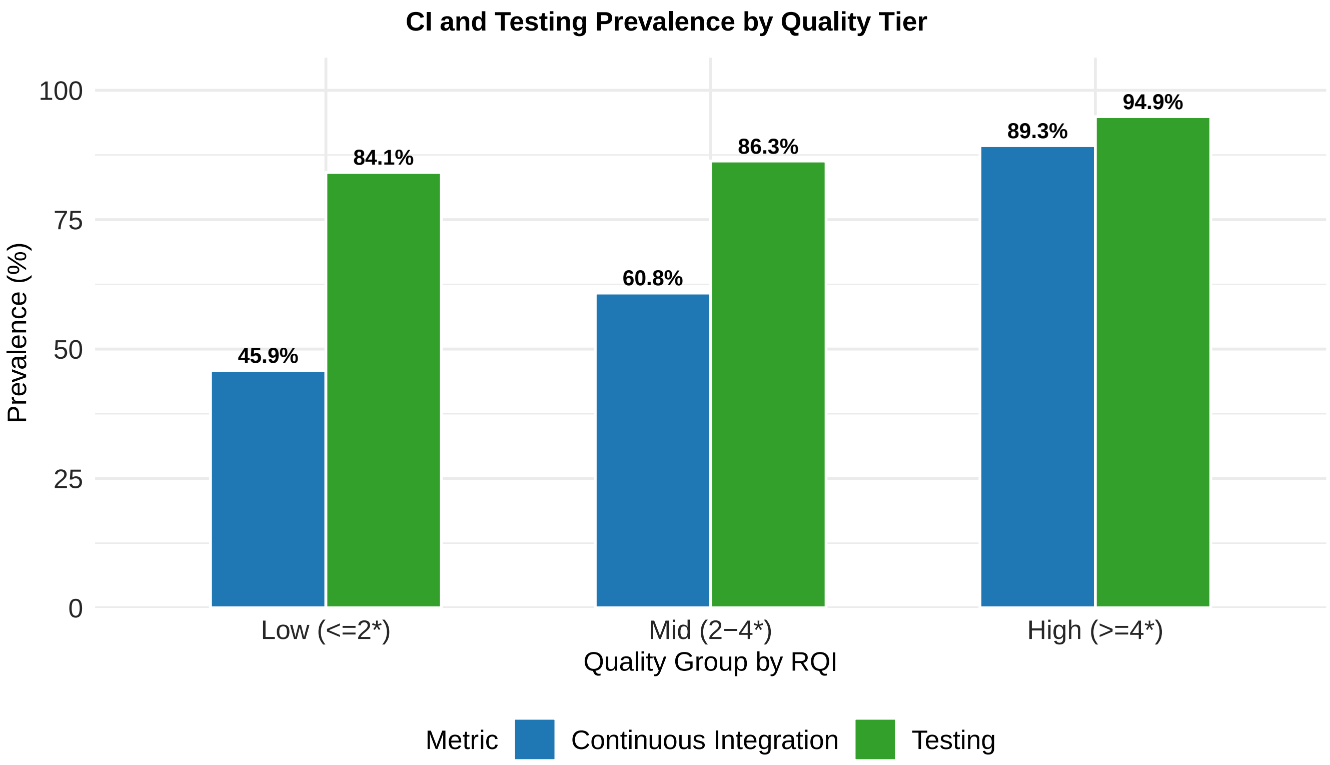


**Table ST15. Holm-adjusted pairwise tests of prevalence between RQI tiers (p values)**

| Comparison | CI | Testing |
|---|---|---|
| 2 vs. 1 | <0.001 | 0.011 |
| 3 vs. 1 | <0.001 | <0.001 |
| 3 vs. 2 | <0.001 | <0.001 |

### 11. CI and Tests by Type

#### 11.1 t-Tests

**Table ST16. Welch two-sample t-tests of RQI by CI and test presence**

| Comparison | Mean 1 | Mean 2 | Difference | t | df | p | CI lower | CI upper |
|---|---|---|---|---|---|---|---|---|
| RQI by CI (1 = no CI, 2 = has CI) | 2.518 | 3.090 | -0.572 | -40.01 | 12676.2 | <0.001 | -0.601 | -0.544 |
| RQI by tests (1 = no tests, 2 = has tests) | 2.628 | 2.914 | -0.286 | -13.95 | 2498.1 | <0.001 | -0.326 | -0.245 |

#### 11.2 Mean RQI by CI and Tests

**Fig S9. Mean RQI by CI and testing presence.** Means ± SE; brackets give Welch t-test p values.

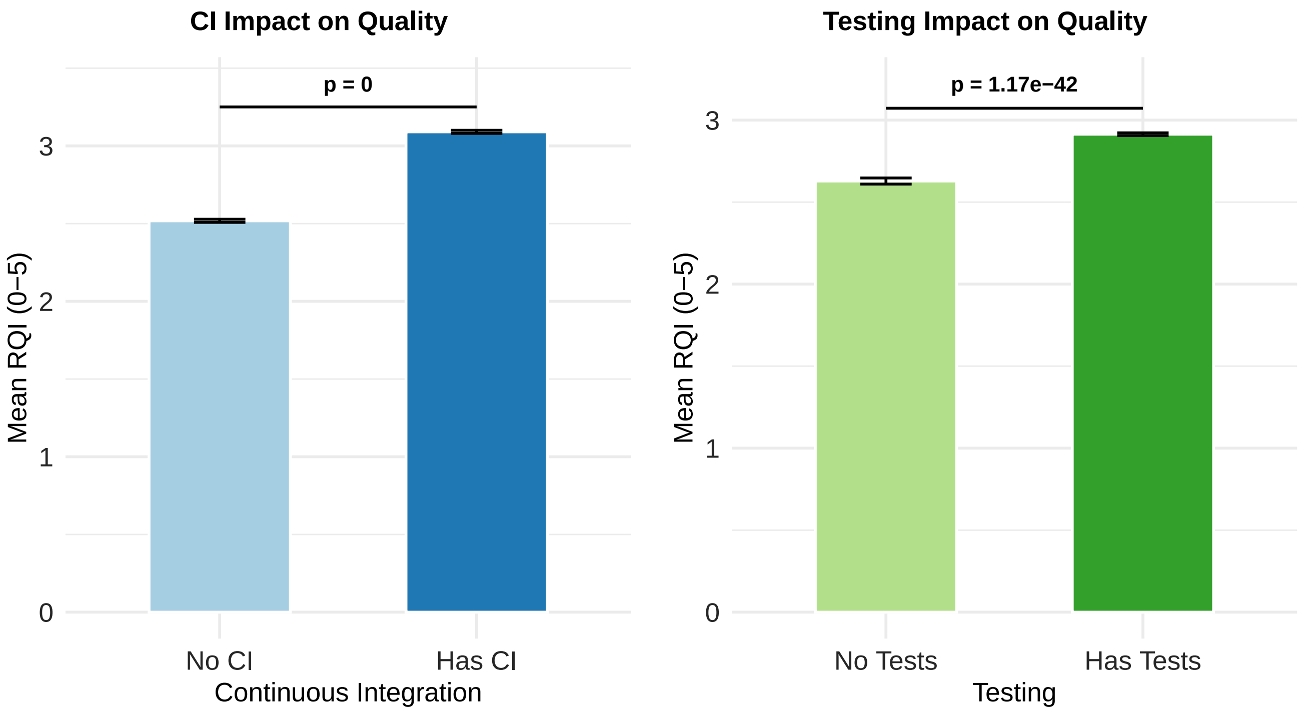


#### 11.3 Within-Type Tests

**Table ST17. Mean RQI with and without CI or tests, by repository type (Welch t-test p values)**

| Type | With CI | No CI | p (CI) | With tests | No tests | p (tests) |
|---|---|---|---|---|---|---|
| astrophysics | 3.16 | 2.45 | <0.001 | 3.02 | 2.45 | 0.003 |
| bioinformatics | 2.92 | 2.53 | <0.001 | 2.82 | 2.52 | <0.001 |
| image_recognition | 2.80 | 2.43 | 0.008 | 2.62 | 2.37 | 0.100 |
| open_source | 3.17 | 2.58 | <0.001 | 3.00 | 2.68 | <0.001 |
| software_engineering | 2.29 | 2.16 | <0.001 | 2.22 | 2.22 | 0.930 |

##### 11.4 CI Effect by Type

**Fig S10. Mean RQI by CI presence, within each type.** Means ± SE; brackets give Welch t-test p values.

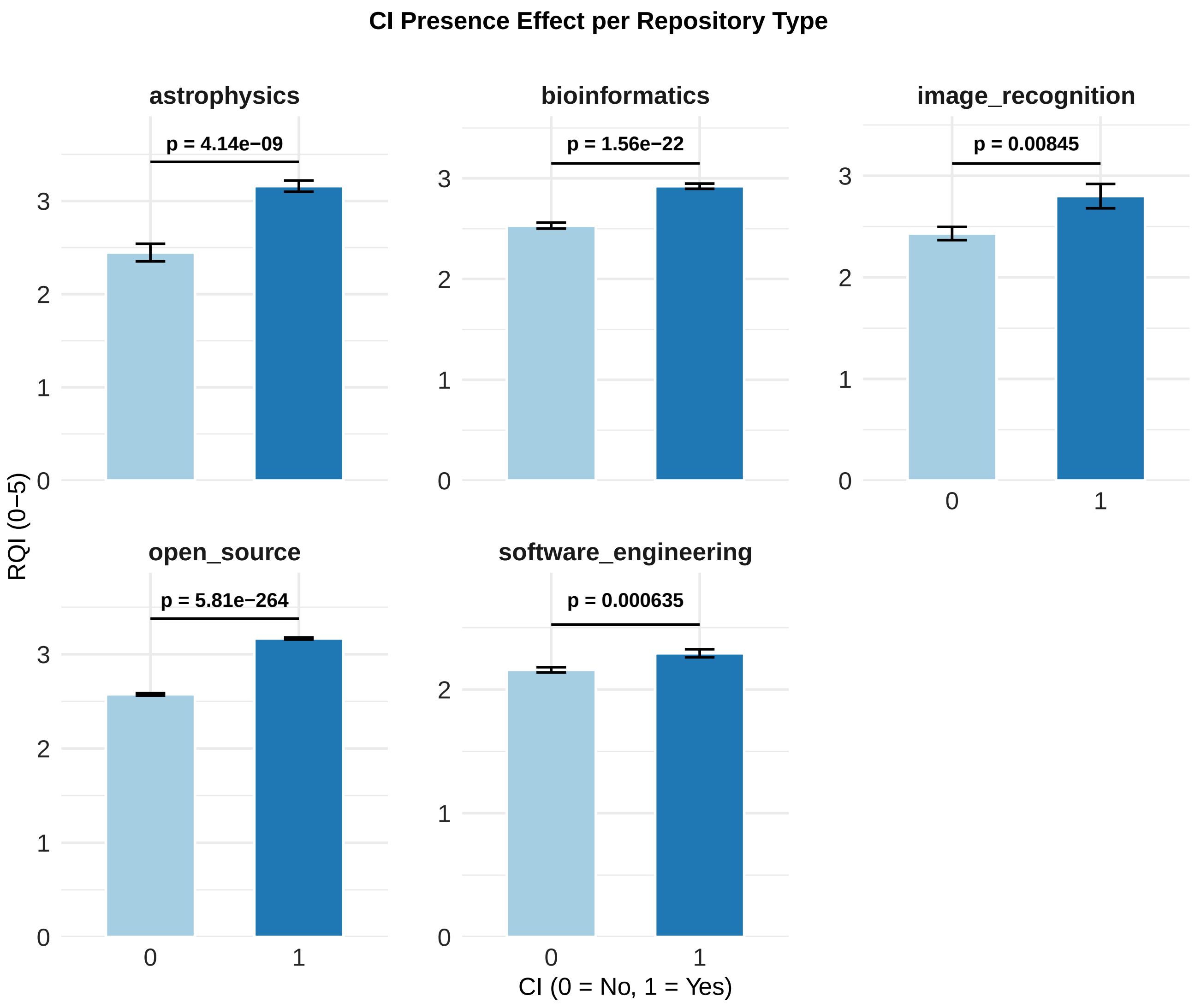

#### 11.5 Testing Effect by Type

**Fig S11. Mean RQI by testing presence, within each type.** Means ± SE; brackets give Welch t-test p values.

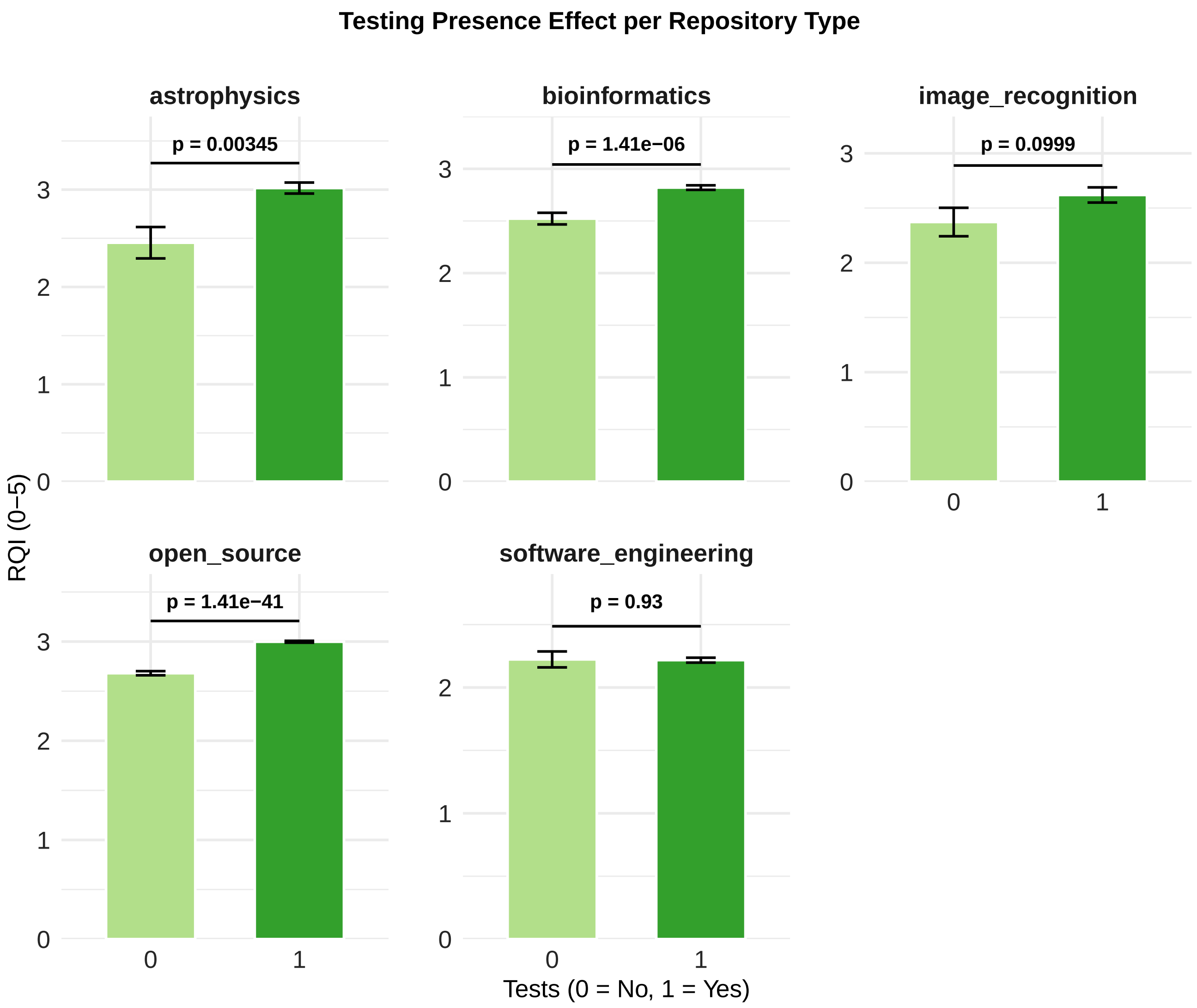

### 12. Languages

#### 12.1 Language List

**Table ST18. Most frequent languages.**

| Language | Repositories |
|---|---|
| JavaScript | 6059 |
| Shell | 5501 |
| HTML | 5499 |
| Python | 5152 |
| CSS | 5104 |
| Dockerfile | 3009 |
| TypeScript | 2436 |
| Makefile | 2057 |
| C++ | 1879 |
| C | 1807 |
| Java | 1661 |
| SCSS | 1106 |
| Batchfile | 938 |
| CMake | 920 |
| C# | 866 |
| PHP | 794 |
| Ruby | 762 |
| Jupyter Notebook | 718 |
| Go | 692 |
| Vue | 444 |
| Kotlin | 443 |
| R | 441 |
| Rust | 410 |
| Objective-C | 405 |
| TeX | 397 |
| PowerShell | 363 |
| Perl | 359 |
| Swift | 359 |
| PLpgSQL | 286 |
| Smarty | 252 |

#### 12.2 Primary Language

Primary language is the one with the most bytes.

**Table ST19. Most frequent primary languages**

| Language | Repositories |
|---|---|
| Python | 2572 |
| JavaScript | 1635 |
| TypeScript | 1505 |
| Java | 1050 |
| C++ | 945 |
| HTML | 624 |
| C# | 611 |
| C | 449 |
| PHP | 443 |
| Go | 413 |
| Jupyter Notebook | 354 |
| Rust | 226 |
| CSS | 212 |
| Kotlin | 175 |
| Shell | 174 |
| Ruby | 166 |
| Dart | 161 |
| R | 159 |
| Vue | 144 |
| Swift | 94 |
| TeX | 72 |
| Julia | 50 |
| Nextflow | 49 |
| Scala | 49 |
| Svelte | 43 |
| SCSS | 42 |
| Objective-C | 39 |
| Perl | 35 |
| Elixir | 34 |
| GDScript | 33 |

#### 12.3 RQI by Language

**Table ST20. Mean RQI by primary language (languages with n  20; top 30 by mean RQI)**

| Language | n | Mean RQI | SD RQI | CI (%) | Tests (%) |
|---|---|---|---|---|---|
| MDX | 28 | 3.52 | 0.79 | 85.7 | 96.4 |
| Rust | 226 | 3.30 | 0.95 | 88.5 | 90.7 |
| Go | 413 | 3.26 | 0.96 | 86.9 | 91.8 |
| Fortran | 27 | 3.22 | 0.84 | 77.8 | 96.3 |
| Pascal | 25 | 3.19 | 0.99 | 56.0 | 72.0 |
| TypeScript | 1505 | 3.15 | 0.96 | 79.8 | 94.0 |
| GDScript | 33 | 3.12 | 1.01 | 54.5 | 90.9 |
| C | 449 | 3.11 | 0.91 | 54.8 | 86.0 |
| Julia | 50 | 3.06 | 0.84 | 92.0 | 98.0 |
| Dart | 161 | 3.01 | 0.88 | 70.2 | 99.4 |
| Kotlin | 175 | 3.00 | 0.96 | 76.6 | 100.0 |
| Shell | 174 | 3.00 | 0.89 | 66.1 | 73.0 |
| C++ | 945 | 2.99 | 0.91 | 60.2 | 89.8 |
| Vue | 144 | 2.97 | 0.95 | 66.7 | 89.6 |
| MATLAB | 28 | 2.96 | 0.95 | 35.7 | 89.3 |
| Python | 2572 | 2.95 | 0.85 | 66.6 | 88.5 |
| Swift | 94 | 2.92 | 0.84 | 61.7 | 98.9 |
| Svelte | 43 | 2.90 | 0.97 | 67.4 | 90.7 |
| Elixir | 34 | 2.90 | 1.01 | 85.3 | 97.1 |
| Lua | 31 | 2.87 | 0.97 | 51.6 | 83.9 |
| C# | 611 | 2.86 | 0.94 | 56.3 | 89.4 |
| Nextflow | 49 | 2.85 | 0.76 | 81.6 | 95.9 |
| PHP | 443 | 2.85 | 0.94 | 57.1 | 91.6 |
| Dockerfile | 22 | 2.82 | 1.10 | 81.8 | 54.5 |
| Jupyter Notebook | 354 | 2.78 | 0.80 | 52.3 | 87.6 |
| R | 159 | 2.75 | 0.77 | 63.5 | 89.3 |
| Perl | 35 | 2.75 | 0.77 | 42.9 | 91.4 |
| Scala | 49 | 2.74 | 0.90 | 89.8 | 100.0 |
| HTML | 624 | 2.66 | 0.84 | 47.0 | 76.8 |
| Java | 1050 | 2.66 | 0.88 | 55.0 | 96.5 |

*Note: JavaScript is not shown because it falls outside the top 30 by mean RQI (rank 33 of 37 languages with n  20). Its values: n = 1635, mean RQI = 2.61, SD = 0.81, CI = 56.9%, tests = 84.5%.*

#### 12.4 RQI by Language, CI and Tests

**Table ST21. Mean RQI by primary language, CI and tests (languages with n  20; top 30 by mean RQI).** Mean RQI (0–5) by presence of tests and CI. JavaScript is not shown because it falls outside the top 30 by mean RQI (rank 33 of 37). Its values: n = 1635, mean RQI = 2.61; no tests = 2.63, has tests = 2.61; no CI = 2.42, has CI = 2.75.

| Language | n | Mean RQI | No tests | Has tests | No CI | Has CI |
|---|---|---|---|---|---|---|
| MDX | 28 | 3.52 | 4.19 | 3.49 | 3.37 | 3.54 |
| Rust | 226 | 3.30 | 2.92 | 3.34 | 2.78 | 3.37 |
| Go | 413 | 3.26 | 3.10 | 3.27 | 2.65 | 3.35 |
| Fortran | 27 | 3.22 | 4.17 | 3.18 | 2.86 | 3.32 |
| Pascal | 25 | 3.19 | 2.92 | 3.30 | 2.91 | 3.42 |
| TypeScript | 1505 | 3.15 | 2.65 | 3.19 | 2.74 | 3.26 |
| GDScript | 33 | 3.12 | 1.96 | 3.24 | 2.75 | 3.43 |
| C | 449 | 3.11 | 2.67 | 3.18 | 2.76 | 3.39 |
| Julia | 50 | 3.06 | 2.96 | 3.06 | 2.80 | 3.08 |
| Dart | 161 | 3.01 | 4.47 | 3.00 | 2.65 | 3.17 |
| Kotlin | 175 | 3.00 | | 3.00 | 2.57 | 3.13 |
| Shell | 174 | 3.00 | 2.87 | 3.04 | 2.64 | 3.18 |
| C++ | 945 | 2.99 | 2.62 | 3.03 | 2.58 | 3.26 |
| Vue | 144 | 2.97 | 2.41 | 3.04 | 2.61 | 3.15 |
| MATLAB | 28 | 2.96 | 2.35 | 3.03 | 2.58 | 3.63 |
| Python | 2572 | 2.95 | 2.71 | 2.98 | 2.66 | 3.10 |
| Swift | 94 | 2.92 | 1.98 | 2.93 | 2.44 | 3.22 |
| Svelte | 43 | 2.90 | 2.24 | 2.97 | 2.36 | 3.16 |
| Elixir | 34 | 2.90 | 1.68 | 2.93 | 2.50 | 2.96 |
| Lua | 31 | 2.87 | 2.74 | 2.90 | 2.54 | 3.18 |
| C# | 611 | 2.86 | 2.48 | 2.91 | 2.45 | 3.18 |
| Nextflow | 49 | 2.85 | 2.05 | 2.88 | 2.38 | 2.95 |
| PHP | 443 | 2.85 | 2.40 | 2.89 | 2.46 | 3.14 |
| Dockerfile | 22 | 2.82 | 2.83 | 2.82 | 2.16 | 2.97 |
| Jupyter Notebook | 354 | 2.78 | 2.63 | 2.80 | 2.52 | 3.02 |
| R | 159 | 2.75 | 2.23 | 2.81 | 2.47 | 2.91 |
| Perl | 35 | 2.75 | 2.35 | 2.78 | 2.61 | 2.93 |
| Scala | 49 | 2.74 | | 2.74 | 2.64 | 2.75 |
| HTML | 624 | 2.66 | 2.52 | 2.70 | 2.40 | 2.94 |
| Java | 1050 | 2.66 | 2.50 | 2.66 | 2.34 | 2.92 |

**Fig S12. Mean RQI by language and CI presence.** Languages with n 20; means ± SE.

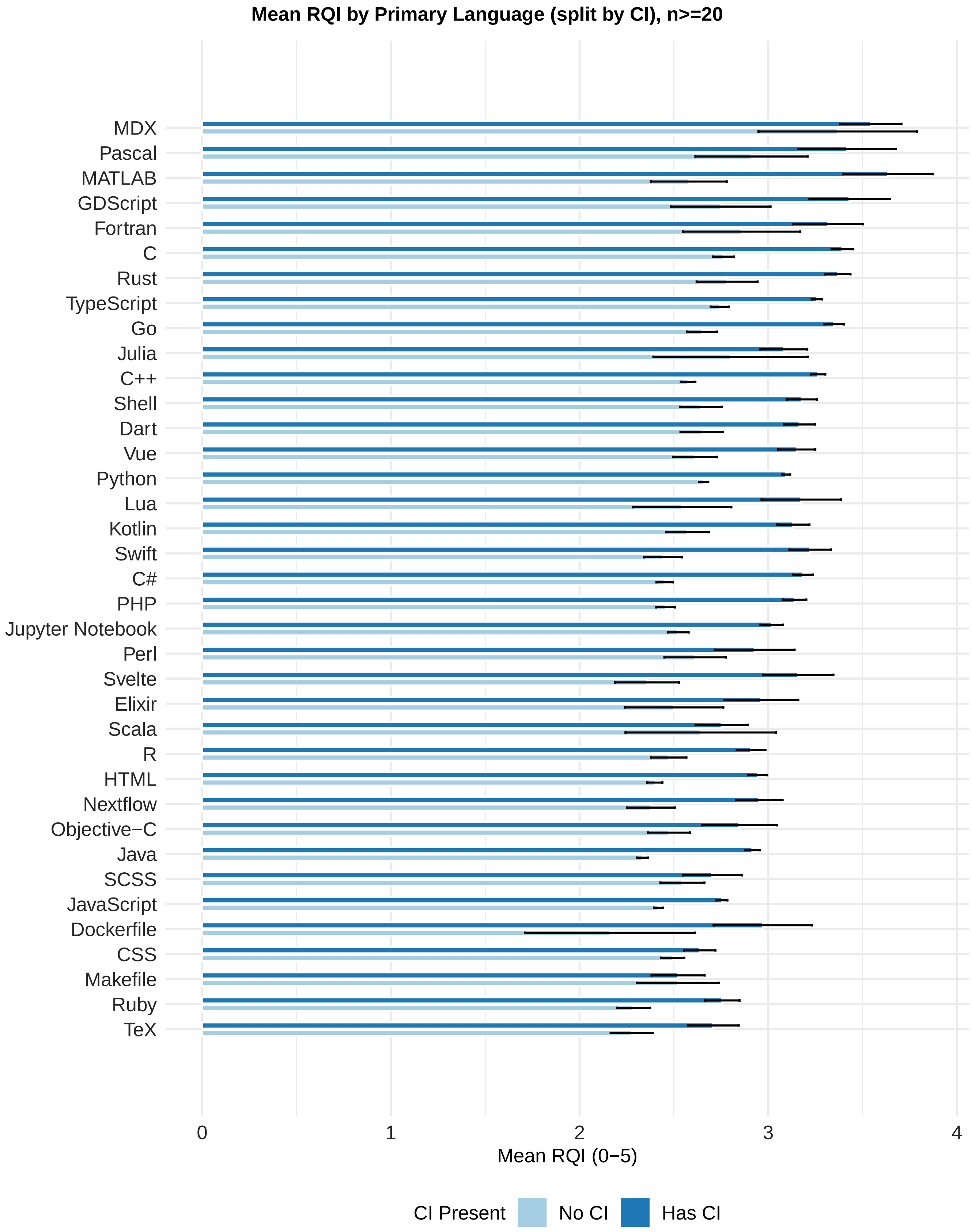

**Fig S13. Mean RQI by language and testing presence.** Languages with n 20; means ± SE.

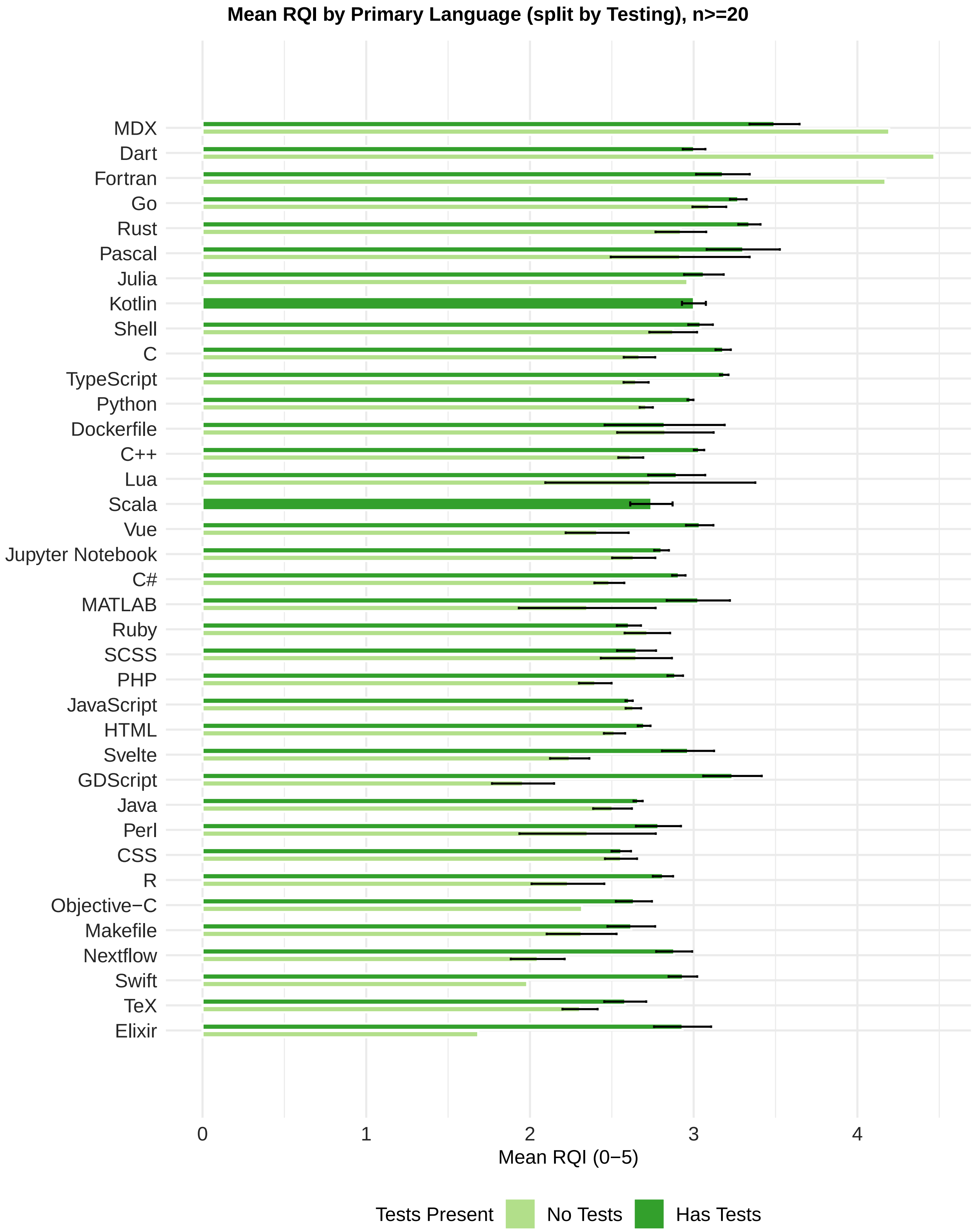

**Fig S14. Mean RQI by language and CI/testing group.** Languages with n 20.

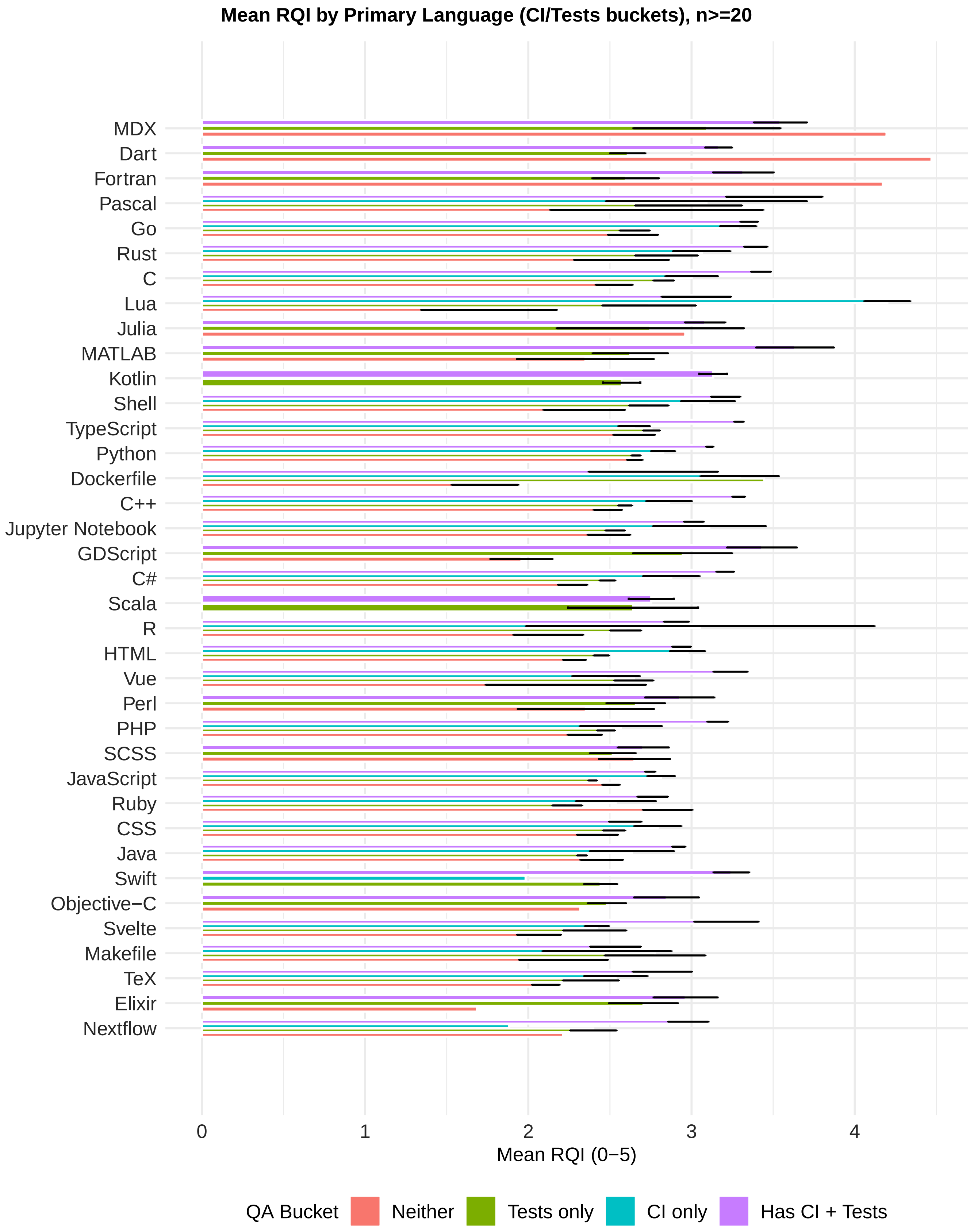

#### 12.5 Bioinformatics Languages

**Table ST22. Most frequent primary languages, bioinformatics**

| Language | Repositories |
|---|---|
| Python | 585 |
| R | 118 |
| C++ | 88 |
| Jupyter Notebook | 80 |
| HTML | 72 |
| JavaScript | 63 |
| Nextflow | 49 |
| Java | 48 |
| Rust | 46 |
| C | 42 |
| Shell | 27 |
| Julia | 26 |
| Perl | 23 |
| TypeScript | 19 |
| TeX | 15 |
| Scala | 14 |
| Go | 12 |
| Ruby | 10 |
| Cython | 6 |
| D | 6 |
| WDL | 6 |
| Groovy | 5 |
| MATLAB | 5 |
| Vue | 4 |
| C# | 3 |
| CSS | 3 |
| Clojure | 3 |
| Kotlin | 3 |
| OCaml | 3 |
| PHP | 3 |

**Table ST23. Most frequent languages, bioinformatics**

| Language | Repositories |
|---|---|
| Python | 978 |
| Shell | 712 |
| HTML | 376 |
| Dockerfile | 350 |
| R | 321 |
| Makefile | 301 |
| JavaScript | 252 |
| CSS | 235 |
| C++ | 177 |
| Jupyter Notebook | 176 |
| C | 159 |
| Perl | 123 |
| TeX | 114 |
| Nextflow | 88 |
| Java | 77 |
| Rust | 59 |
| CMake | 56 |
| Batchfile | 49 |
| SCSS | 45 |
| TypeScript | 42 |
| Cython | 34 |
| Roff | 31 |
| Ruby | 31 |
| Julia | 30 |
| Groovy | 25 |
| WDL | 23 |
| Scala | 22 |
| Go | 19 |
| M4 | 18 |
| Awk | 16 |

**Table ST24. Mean RQI by primary language, bioinformatics (languages with n  10; top 30 by mean RQI)**

| Language | n | Mean RQI | SD RQI | CI (%) | Tests (%) |
|---|---|---|---|---|---|
| Go | 12 | 3.15 | 0.79 | 91.7 | 100.0 |
| Rust | 46 | 3.06 | 0.68 | 84.8 | 87.0 |
| C | 42 | 3.04 | 0.74 | 50.0 | 81.0 |
| TypeScript | 19 | 2.96 | 0.80 | 89.5 | 94.7 |
| Java | 48 | 2.88 | 0.92 | 62.5 | 97.9 |
| Julia | 26 | 2.87 | 0.86 | 84.6 | 96.2 |
| Python | 585 | 2.85 | 0.76 | 73.3 | 92.1 |
| Nextflow | 49 | 2.85 | 0.76 | 81.6 | 95.9 |
| C++ | 88 | 2.80 | 0.66 | 63.6 | 92.0 |
| JavaScript | 63 | 2.77 | 0.71 | 65.1 | 92.1 |
| R | 118 | 2.76 | 0.75 | 62.7 | 88.1 |
| Shell | 27 | 2.74 | 0.88 | 37.0 | 74.1 |
| Ruby | 10 | 2.65 | 0.99 | 80.0 | 100.0 |
| Jupyter Notebook | 80 | 2.59 | 0.71 | 42.5 | 83.8 |
| Perl | 23 | 2.58 | 0.65 | 34.8 | 91.3 |
| Scala | 14 | 2.57 | 1.01 | 100.0 | 100.0 |
| HTML | 72 | 2.56 | 0.86 | 51.4 | 83.3 |
| TeX | 15 | 2.39 | 0.66 | 53.3 | 66.7 |

**Table ST25. Mean RQI by primary language, CI and tests, bioinformatics (languages with n  10; top 30 by mean RQI).** Mean RQI (0–5) by presence of tests and CI.

| Language | n | Mean RQI | No tests | Has tests | No CI | Has CI |
|---|---|---|---|---|---|---|
| Go | 12 | 3.15 | | 3.15 | 3.76 | 3.10 |
| Rust | 46 | 3.06 | 2.85 | 3.09 | 2.94 | 3.08 |
| C | 42 | 3.04 | 2.85 | 3.08 | 2.86 | 3.22 |
| TypeScript | 19 | 2.96 | 2.27 | 2.99 | 3.29 | 2.92 |
| Java | 48 | 2.88 | 2.23 | 2.90 | 2.64 | 3.03 |
| Julia | 26 | 2.87 | 2.96 | 2.86 | 2.80 | 2.88 |
| Python | 585 | 2.85 | 2.62 | 2.87 | 2.58 | 2.95 |
| Nextflow | 49 | 2.85 | 2.05 | 2.88 | 2.38 | 2.95 |
| C++ | 88 | 2.80 | 2.92 | 2.79 | 2.55 | 2.94 |
| JavaScript | 63 | 2.77 | 2.46 | 2.79 | 2.67 | 2.82 |
| R | 118 | 2.76 | 2.23 | 2.83 | 2.55 | 2.88 |
| Shell | 27 | 2.74 | 2.80 | 2.71 | 2.59 | 2.99 |
| Ruby | 10 | 2.65 | | 2.65 | 2.32 | 2.74 |
| Jupyter Notebook | 80 | 2.59 | 2.35 | 2.64 | 2.51 | 2.70 |
| Perl | 23 | 2.58 | 2.14 | 2.62 | 2.55 | 2.63 |
| Scala | 14 | 2.57 | | 2.57 | | 2.57 |
| HTML | 72 | 2.56 | 2.37 | 2.60 | 2.23 | 2.87 |
| TeX | 15 | 2.39 | 2.34 | 2.42 | 2.47 | 2.32 |

**Fig S15. Mean RQI by language, bioinformatics.** Languages with n  10.

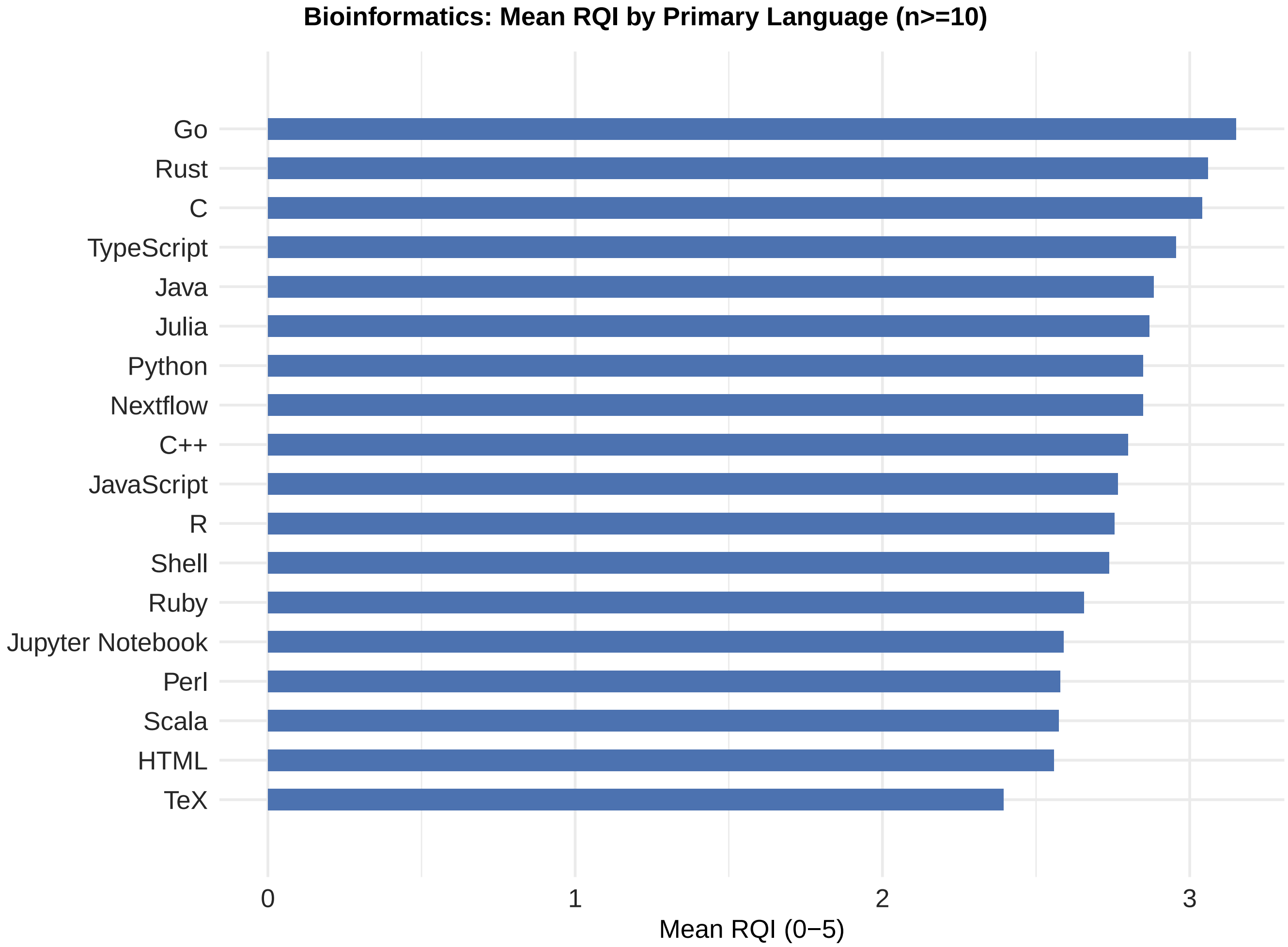

**Fig S16. Mean RQI by language and CI presence, bioinformatics.** Means ± SE.

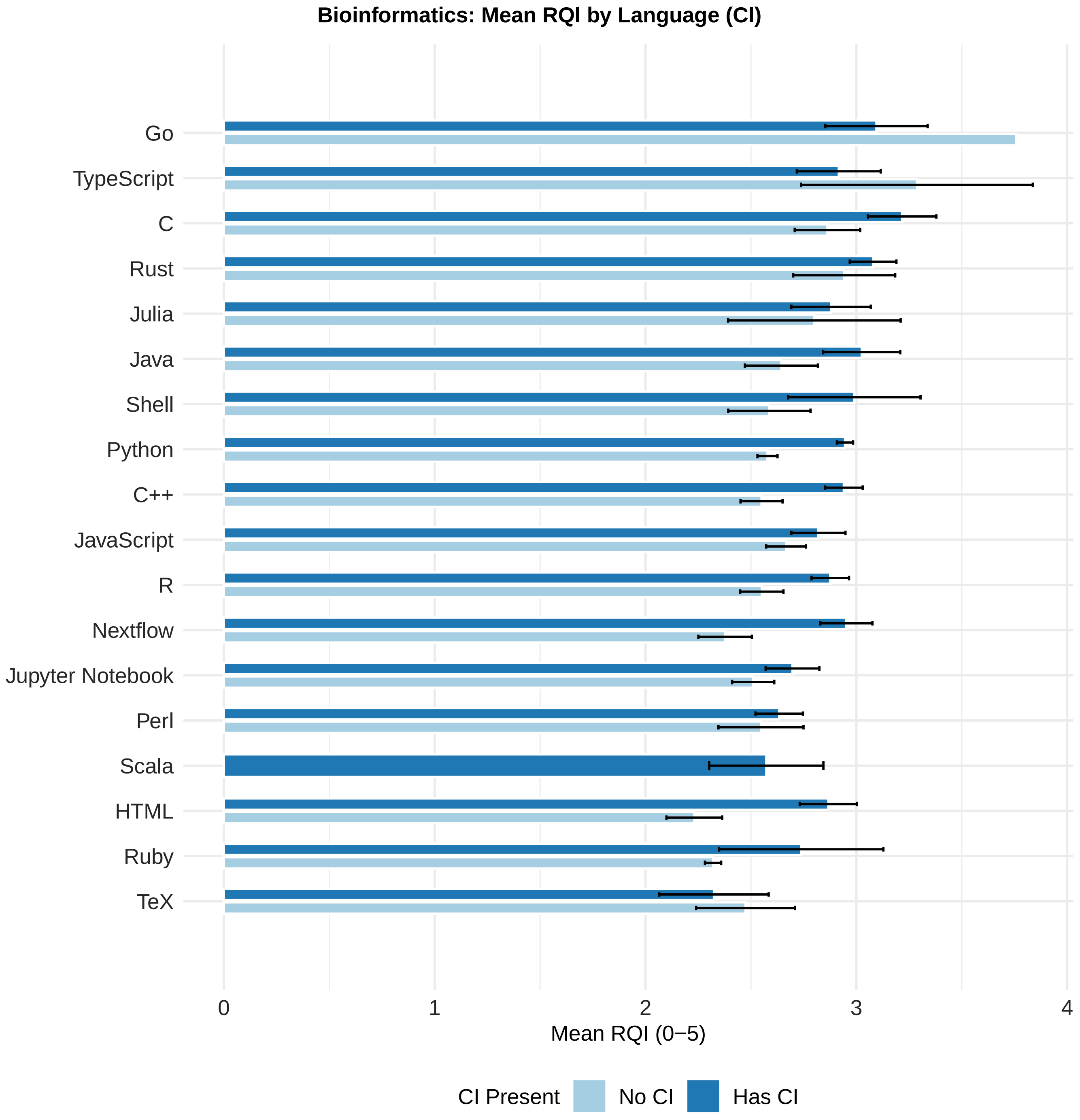

**Fig S17. Mean RQI by language and testing presence, bioinformatics.** Means ± SE.

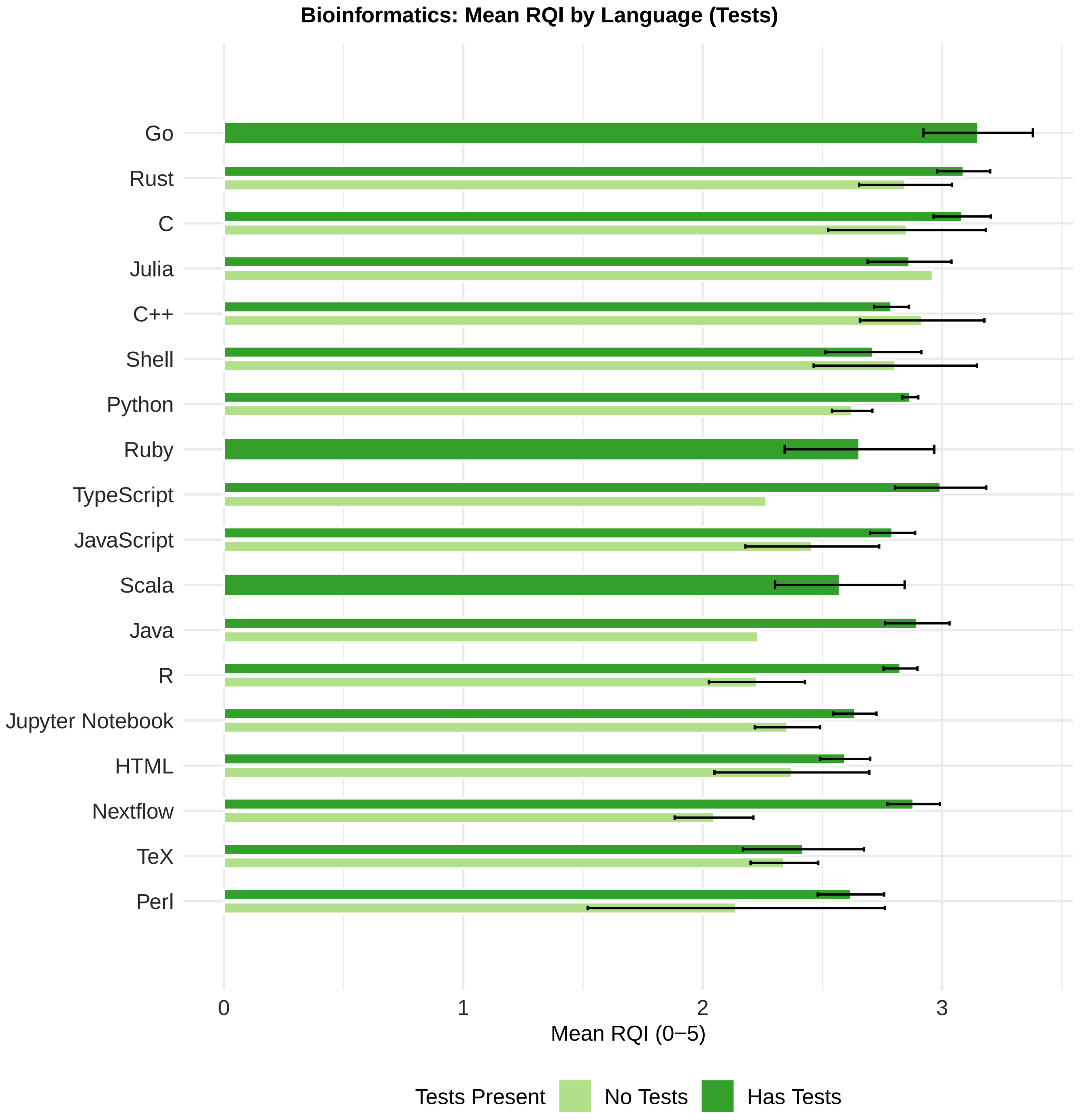

#### 13. Repository Age

Repositories are grouped into age cohorts; mean RQI is compared by CI and testing.

##### 13.1 Age and CI

**Fig 3A (main text). Mean RQI by age cohort and CI status.** Age is years from first commit to data collection (March 2026); means ± SE.

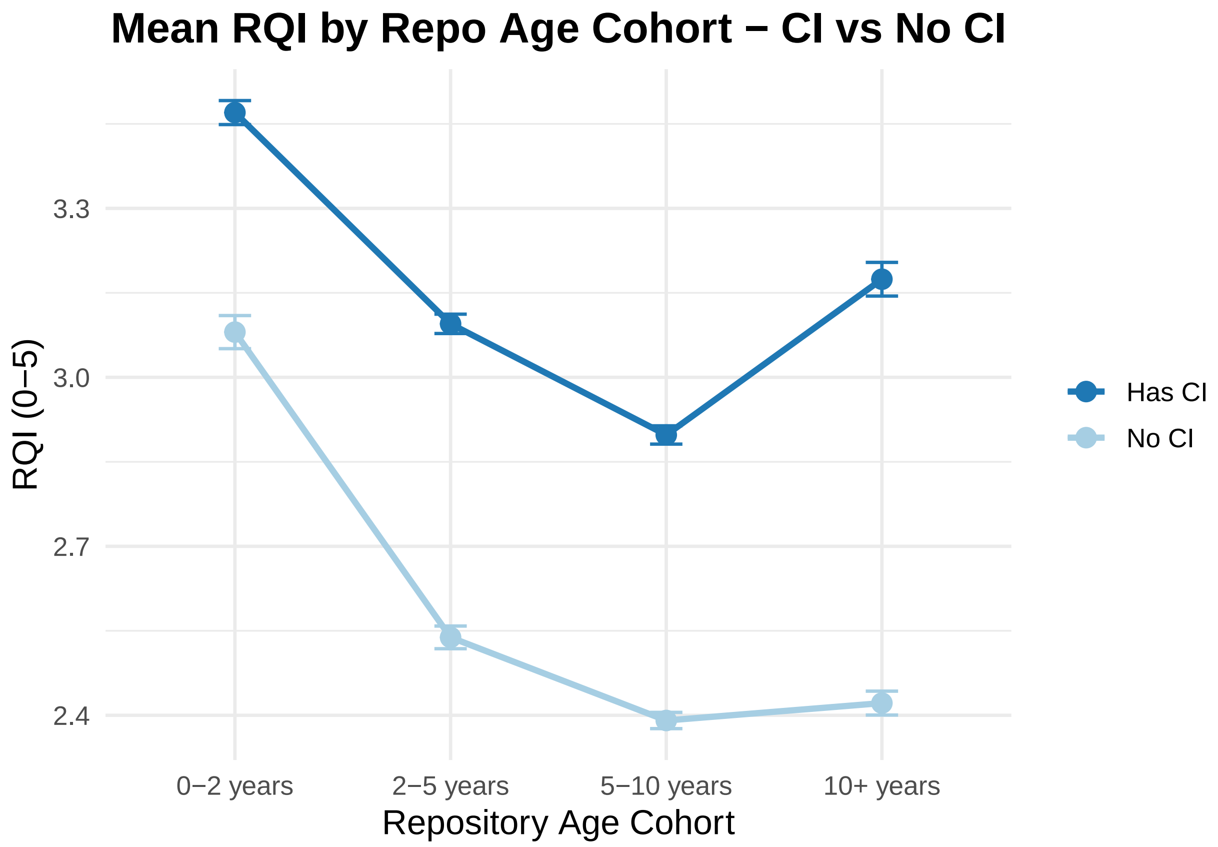

##### 13.2 Age and Testing

**Fig 3B (main text). Mean RQI by age cohort and testing status.** Means ± SE.

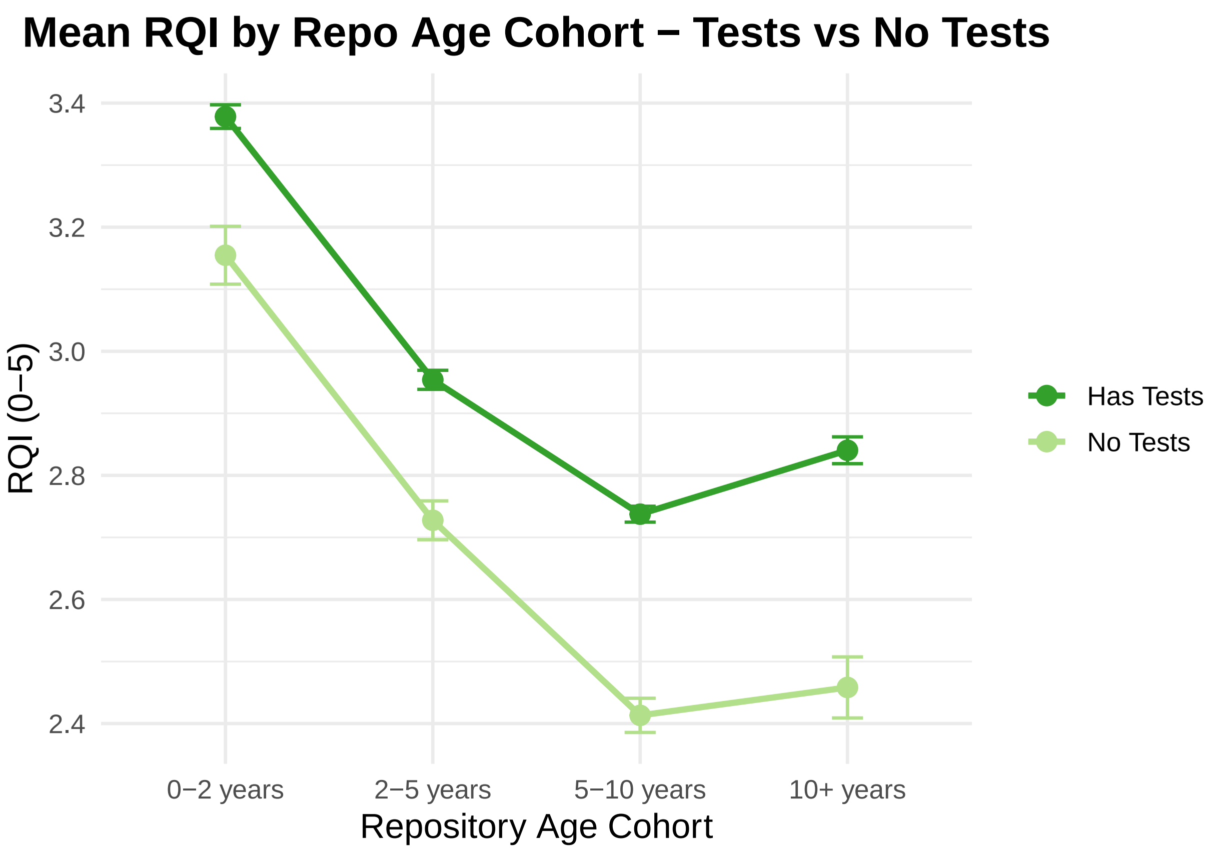

##### 13.3 Age, CI and Testing (Combined)

**Fig 3 (main text). Mean RQI by age cohort, by CI (a) and testing (b).** Age is years from first commit to data collection (March 2026); means ± SE.

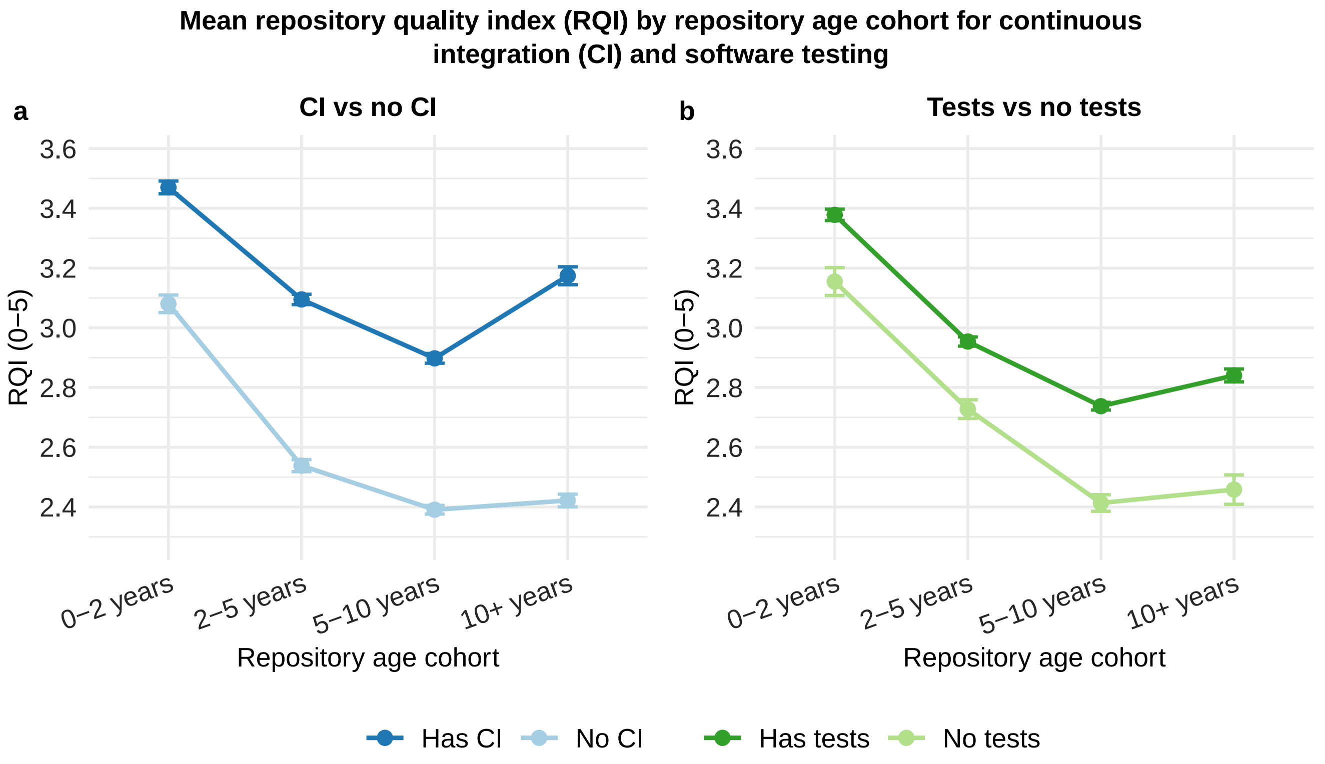

#### 13.4 Cohort Gaps

RQI gaps (with minus without CI or tests) and Welch t-test p-values by cohort.

**Table ST26. Mean RQI by age cohort, with and without CI (Welch t-test p values)**

| Age cohort | No CI | Has CI | Gap | p |
|---|---|---|---|---|
| 0–2 years | 3.08 | 3.47 | 0.39 | <0.001 |
| 2–5 years | 2.54 | 3.09 | 0.56 | <0.001 |
| 5–10 years | 2.39 | 2.90 | 0.51 | <0.001 |
| 10+ years | 2.42 | 3.17 | 0.75 | <0.001 |

**Table ST27. Mean RQI by age cohort, with and without tests (Welch t-test p values)**

| Age cohort | No tests | Has tests | Gap | p |
|---|---|---|---|---|
| 0–2 years | 3.15 | 3.38 | 0.22 | <0.001 |
| 2–5 years | 2.73 | 2.95 | 0.23 | <0.001 |
| 5–10 years | 2.41 | 2.74 | 0.32 | <0.001 |
| 10+ years | 2.46 | 2.84 | 0.38 | <0.001 |

### 14. Citations and RQI

#### 14.1 Median Citations, Published Repos

**Fig 5 (main text). Median citations by RQI bin, published repositories.** Repositories with at least one citation; color shows RQI (red = low, green = high). Brackets give exact p-values for significant Wilcoxon comparisons.

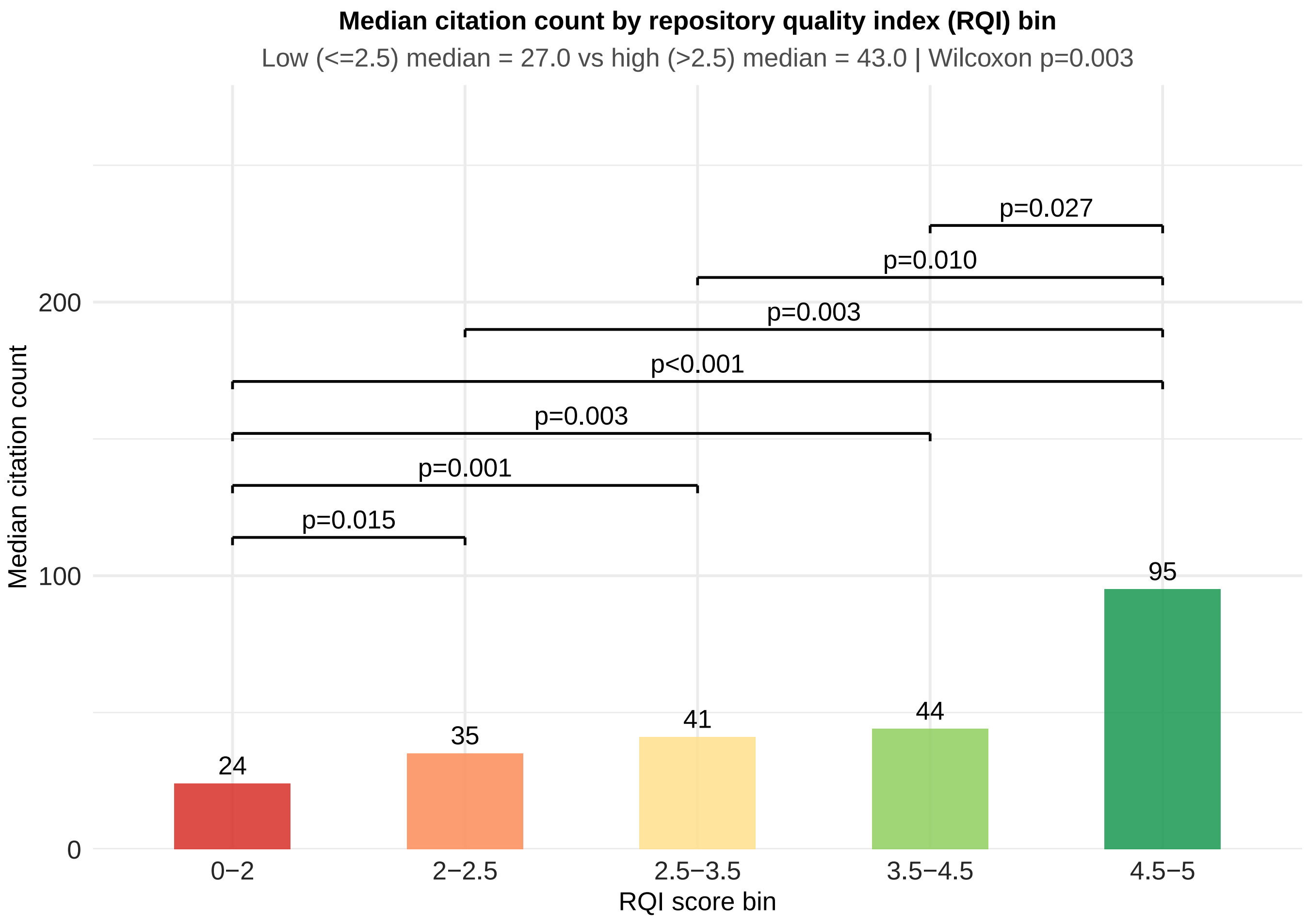

### 15. Publication Status

#### 15.1 RQI Components

**Fig 4 (main text), panel a. RQI and components by publication status, bioinformatics.** Combined with panel b in Section 16.4.

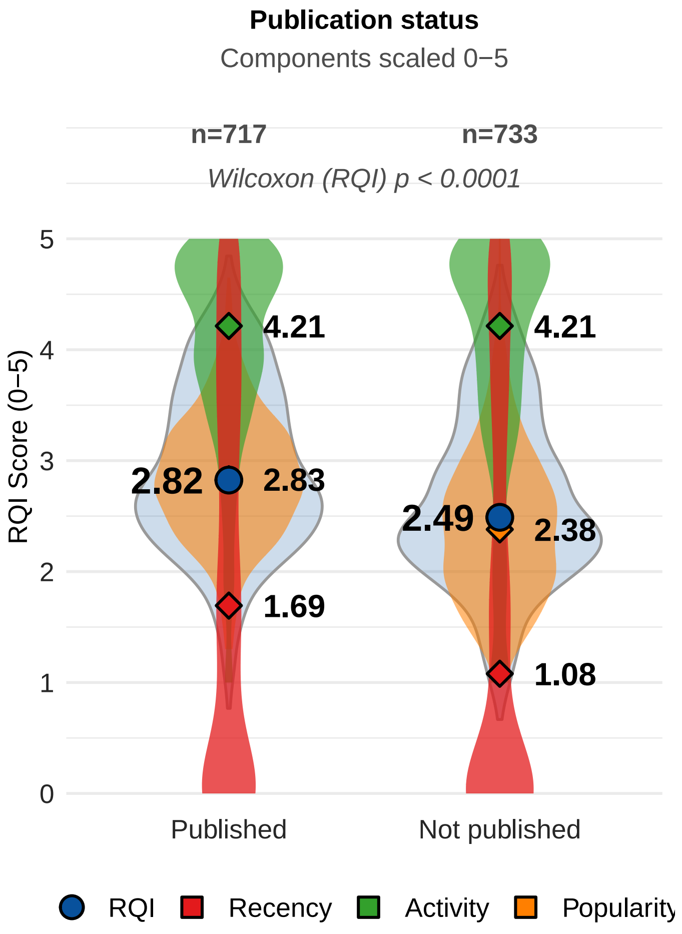

### 16. Funding

#### 16.1 Data

**Table ST28. Funding data overview**

| Statistic | Count |
|---|---|
| Repositories | 1257 |
| Grant funded | 674 |
| Not grant funded | 583 |
| With developer award and RQI | 175 |

#### 16.2 RQI Components (All Domains)

**Fig S18. RQI and components by funding status, all domains.**

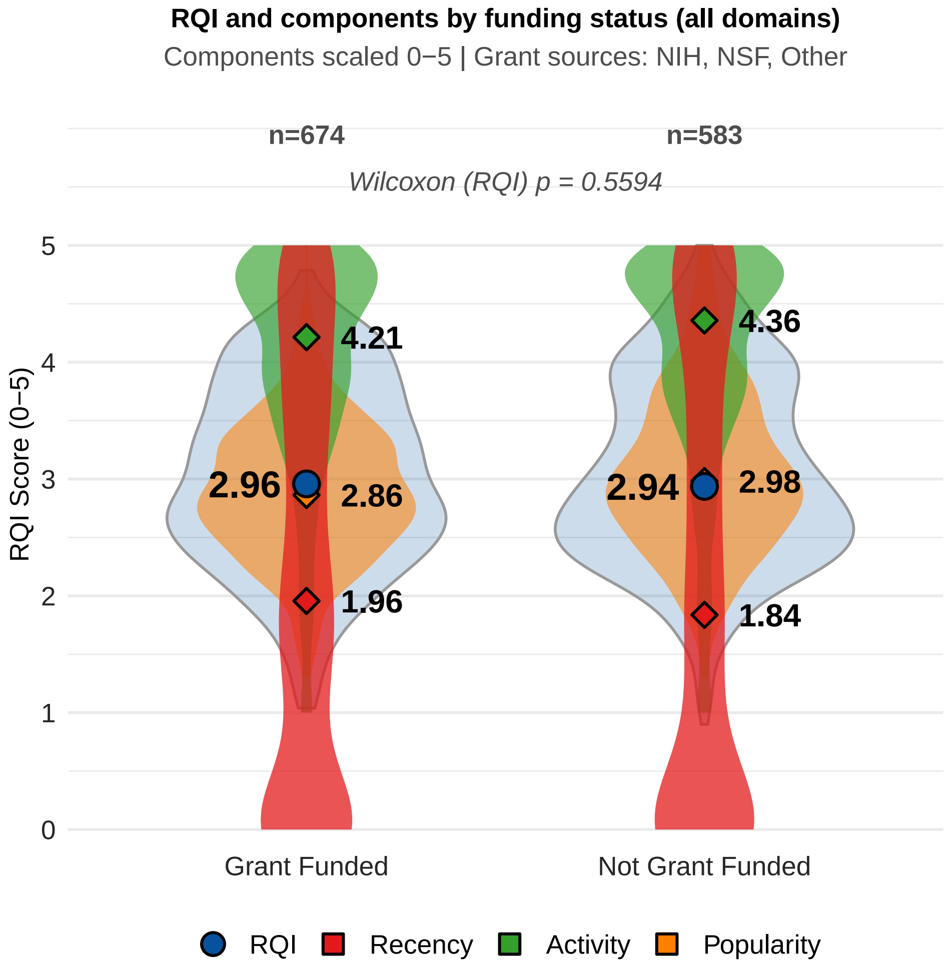

#### 16.3 Test Statistics (All Domains)

**Table ST29. Wilcoxon rank-sum test of RQI by grant funding status (all domains)**

| Comparison | n 1 | n 2 | Median 1 | Median 2 | W | p | Rank-biserial r |
|---|---|---|---|---|---|---|---|
| RQI (1 = Grant Funded, 2 = Not Grant Funded) | 674 | 583 | 2.96 | 2.94 | 192724 | 0.559 | 0.019 |

#### 16.4 RQI Components (Bioinformatics)

**Fig 4 (main text), panel b. RQI and components by funding status, bioinformatics.**

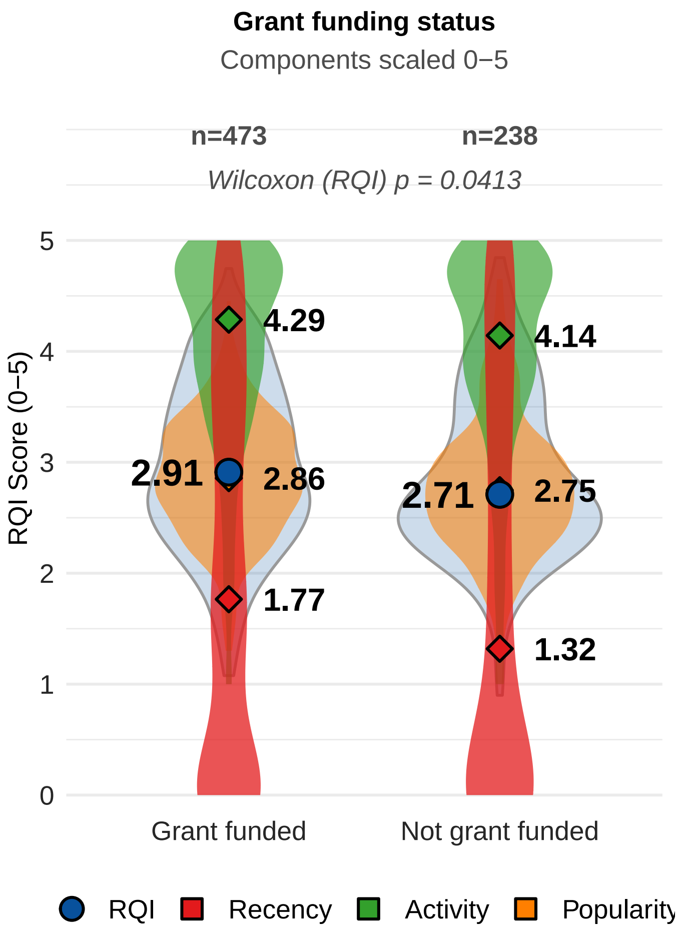


*Grant sources:* NIH, NSF, Other.

**Table ST30. Wilcoxon rank-sum test of RQI by grant funding status (bioinformatics)**

| Comparison | n 1 | n 2 | Median 1 | Median 2 | W | p | Rank-biserial r |
|---|---|---|---|---|---|---|---|
| RQI (1 = Grant Funded, 2 = Not Grant Funded) | 473 | 238 | 2.91 | 2.71 | 61561.5 | 0.041 | -0.094 |

**Fig 4 (main text). RQI and components by publication (a) and grant funding (b), bioinformatics.** Violins show RQI and its components; diamonds mark medians; Wilcoxon results annotated.

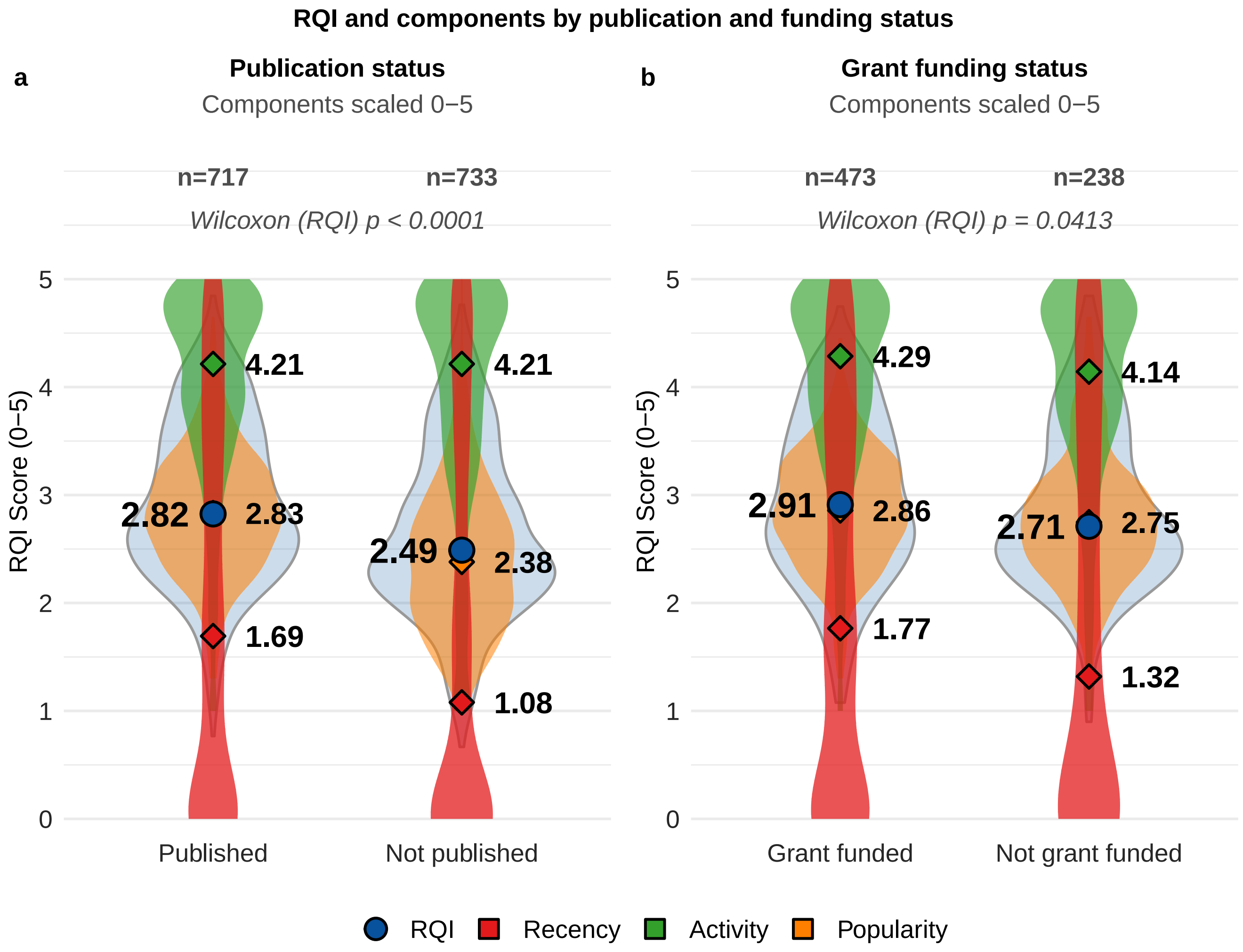

### 17. Citation Models

#### 17.1 Transformations

**Fig S19. Citation transformation diagnostics.** Raw, log, and square-root histograms, residual plots, and Box–Cox profile.

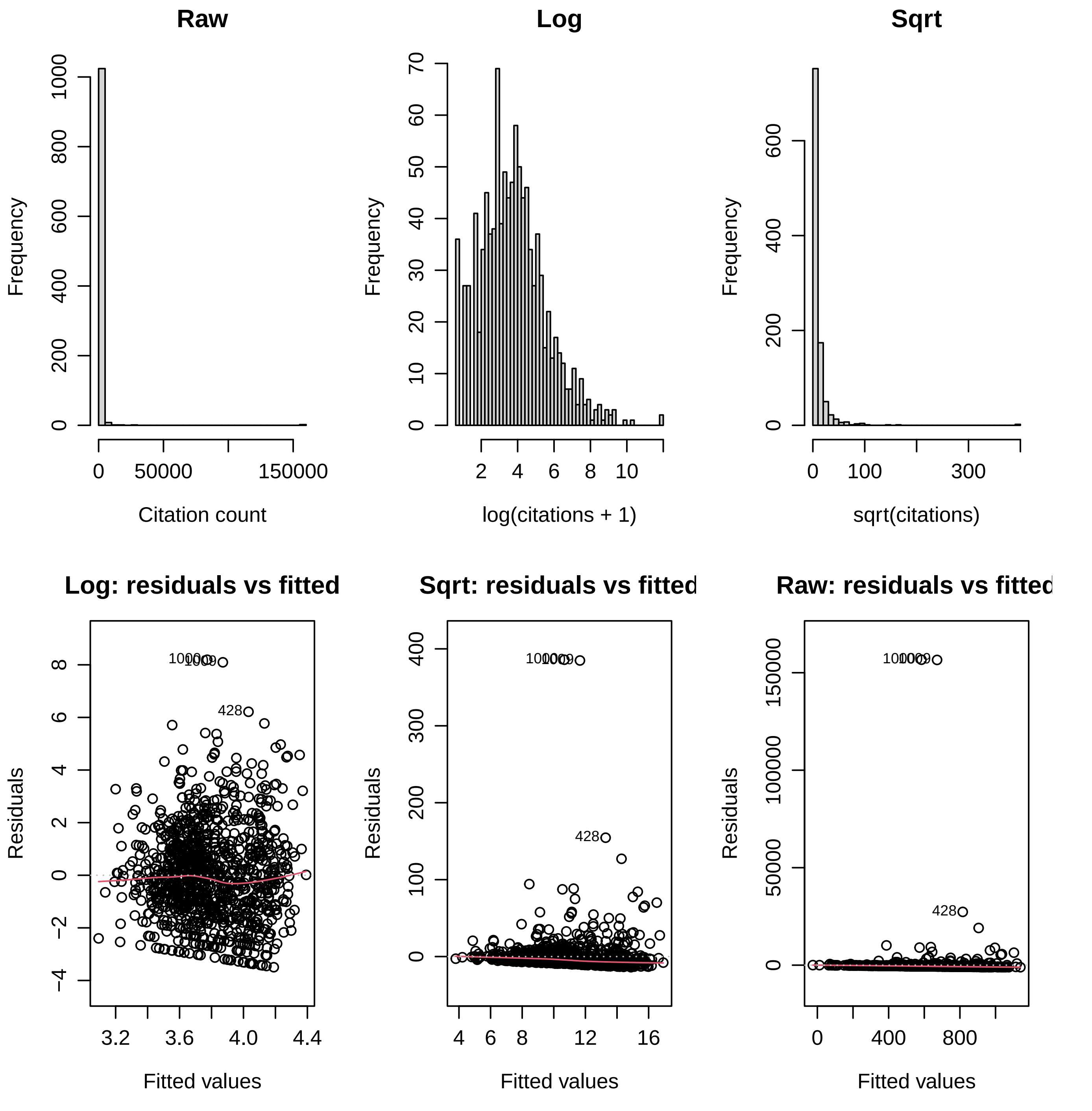

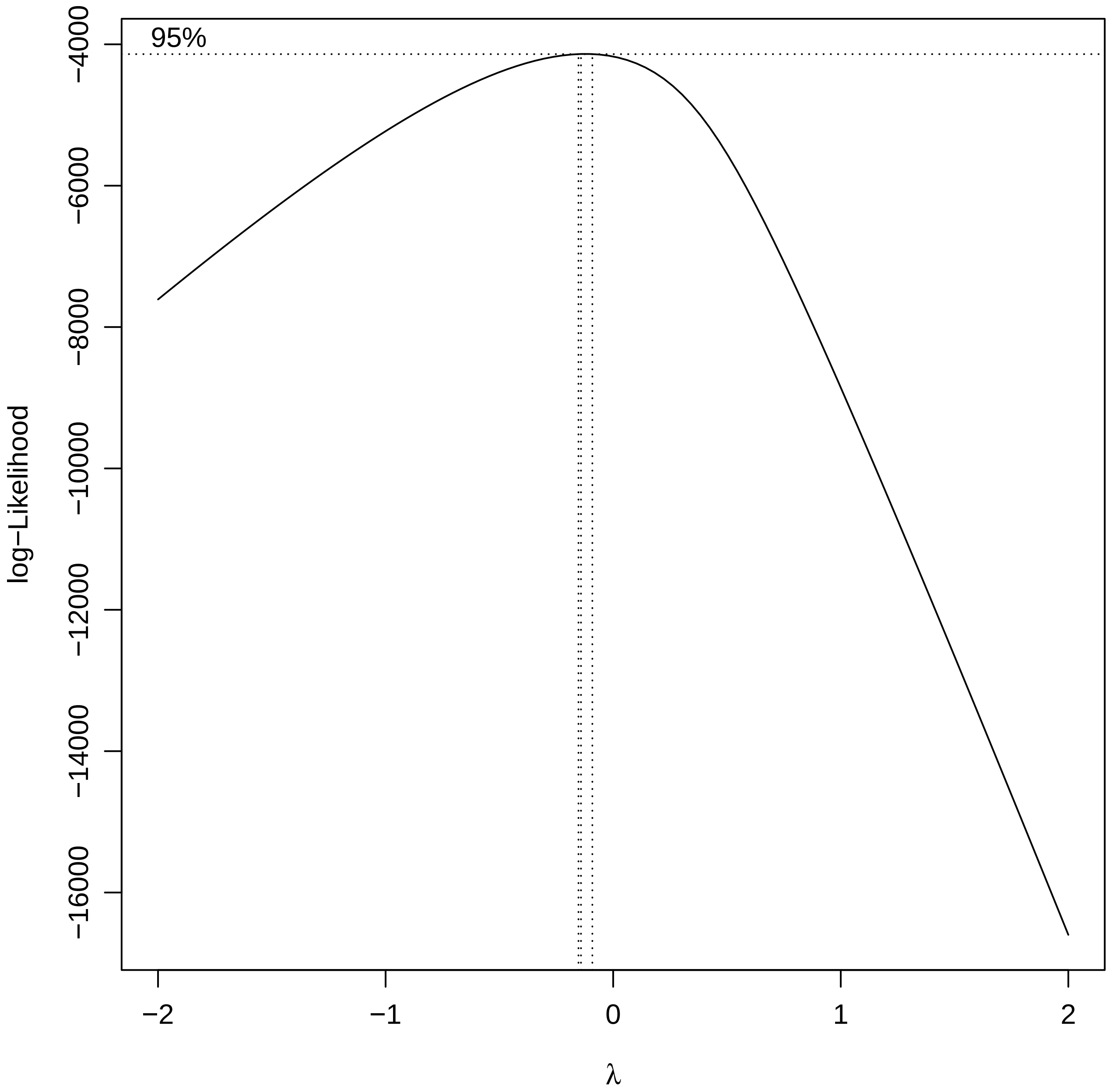


**Table ST31. Citation transformations: model R-squared and Box-Cox lambda**

| R-squared (log) | R-squared (sqrt) | R-squared (raw) | Box-Cox lambda |
|---:|---:|---:|---:|
| 0.019 | 0.013 | 0.001 | -0.141 |

#### 17.2 Citations vs RQI

**Fig S20. Citations vs RQI, published repositories.** OLS fit of log(citations + 1) with 95% CI.

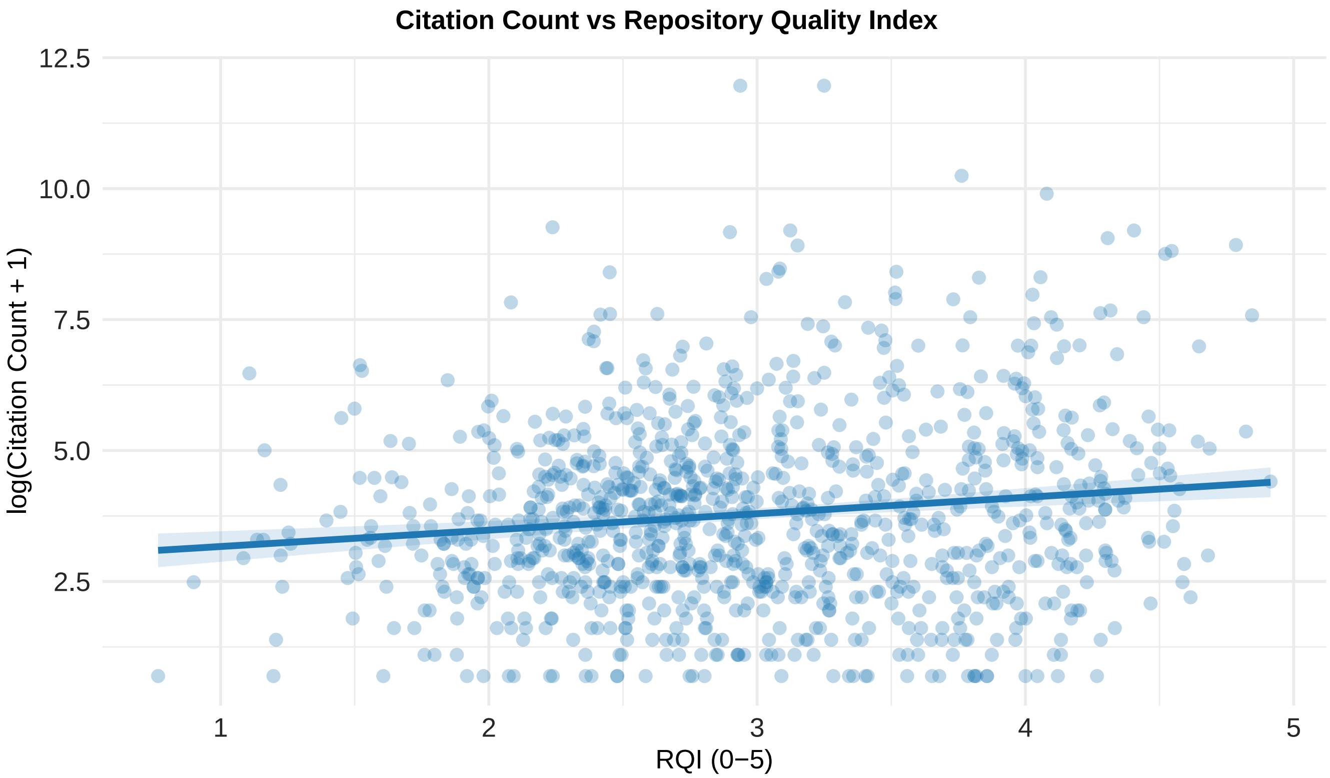


**Table ST32. Linear model of log(citations + 1) on RQI (published repositories)**

| Term | Estimate | Std. Error | t | p | CI lower | CI upper |
|---|---|---|---|---|---|---|
| Intercept | 2.853 | 0.215 | 13.279 | $<0.001$ | 2.432 | 3.275 |
| RQI | 0.313 | 0.070 | 4.499 | $<0.001$ | 0.177 | 0.450 |

**Table ST33. Model fit (log(citations + 1) on RQI)**

| n | R-squared | Adj. R-squared | Residual SE | Residual df | F | F df | p |
|---|---|---|---|---|---|---|---|
| 1037 | 0.019 | 0.018 | 1.737 | 1035 | 20.241 | 1, 1035 | $<0.001$ |
